# Single-cell and spatial transcriptomics reveals the human liver immunological landscape and myeloid dysfunction in PSC

**DOI:** 10.1101/2023.07.28.550550

**Authors:** Tallulah S. Andrews, Diana Nakib, Catia Perciani, Xue Zhong Ma, Lewis Liu, Erin Winter, Damra Camat, Sai Chung, Justin Manuel, Shantel Mangroo, Bettina Hansen, Bal Arpinder, Cornelia Thoeni, Blayne Sayed, Jordan Feld, Adam Gehring, Aliya Gulamhusein, Gideon M Hirschfield, Amanda Riciutto, Gary D. Bader, Ian D. McGilvray, Sonya MacParland

## Abstract

**Background:** Primary sclerosing cholangitis (PSC) is an immune-mediated cholestatic liver disease characterized by bile retention, biliary tree destruction, and progressive fibrosis leading to end stage liver disease and transplantation. There is an unmet need to understand the cellular composition of the PSC liver and how it underlies disease pathogenesis. As such, we generated a comprehensive atlas of the PSC liver and a reference healthy liver dataset using multiple multi-omic modalities and functional validation.

**Methods:** In this work, we employed single-cell (12,000 cells), single-nuclei (23,000 nuclei), and spatial transcriptomics (1 sample by 10x Visium and 3 samples with multi-region profiling by Nanostring GeoMx DSP) to profile the cellular ecosystem in 5 patients with PSC. Transcriptomic profiles were compared to 100k single cell transcriptomes and spatial transcriptomics controls from 24 healthy neurologically deceased donor (NDD) livers. Flow cytometry and intracellular cytokine staining was performed to validate PSC-specific differences in immune phenotype and function.

**Results:** PSC explants with cirrhosis of the liver parenchyma and prominent periductal fibrosis were associated with a unique population of hepatocytes which transformed to a cholangiocyte-like phenotype. These hepatocytes were surrounded by diverse immune cell populations, including monocyte-like macrophages, liver-resident and circulating natural killer (NK) cells. Inflamed cholangiocytes, fibrosis-resident hepatic stellate cells, and endothelial cells released cytokines that recruited CD4+T-cells, dendritic cells, and neutrophils to the PSC liver. Tissue-resident macrophages, by contrast, were reduced in number and exhibited a dysfunctional inflammatory response to LPS and IFN-Ɣ stimulation.

**Conclusions:** We present the first comprehensive atlas of the PSC liver and demonstrate hyper-activation and exhaustion-like phenotypes of myeloid cells and markers of chronic cytokine expression in late-stage PSC lesions.

**Lay Summary:** Primary sclerosing cholangitis (PSC) is a rare liver disease characterized by chronic inflammation and irreparable damage to the bile ducts. Due to a limited understanding of the underlying pathogenesis of disease, there remains a paucity of treatment options. As such, we sequenced healthy and diseased livers to compare the activity, interactions, and localization of immune and non-immune cells. This revealed that outside PSC scar regions, hepatocytes are transitioning to bile duct cells, whereas within the scars, there is an accumulation of immune cells. Of these cells, macrophages that typically contribute to tissue repair were enriched in immunoregulatory genes and were less responsive to stimulation. These cells are likely involved in maintaining hepatic inflammation and could be targeted in novel therapeutic development.

## Introduction

Organ transcriptomic mapping efforts such as the human cell atlas have the potential to revolutionize our understanding of organ biology (1). Recent atlasing efforts have revealed a broad diversity of parenchymal, progenitor, tissue resident and transient circulating immune cells within the liver(2–4). Changes in the frequency and characteristics of these populations have also been observed in atlases of acute liver disease, chronic fibrosis, and hepatic cancers (5–9). Understanding the healthy liver by building a comprehensive reference map with an inventory of cells and their respective spatial localization is crucial in elucidating the transcriptomic and phenotypic changes, rare cell types, and cell-cell interactions that are underlying disease development.

Primary sclerosing cholangitis (PSC) is an immune-mediated cholestatic liver disease that is characterized by the retention of bile, destruction of the biliary tree and development of fibrosis(10). Given the rarity of the disease and the timeline from diagnosis to end-stage liver disease, few studies have attempted to map the PSC liver and characterize the cellular landscape. Previous maps of PSC have mainly considered sorted populations of immune cells(11,12). These and other studies using bulk tissue expression have variously implicated CD4+ T cells(11), neutrophils(13), dendritic cells(12), antibody-producing B cells(14,15), and macrophages(16–21) in the development of disease. However, the narrow scope and low cellular resolution of these studies leave many uncertainties in the underlying cause and consequences of PSC inflammation preventing the development of targeted treatments.

We present the first unbiased atlas of PSC using 5’ single-cell, 3’ single-nucleus and spatial transcriptomics, and a complementary 100k cell reference healthy liver map. These maps revealed 7 PSC-associated cell subtypes including cytotoxic T-cells, dendritic cells, and neutrophils, as well as specific cell-cell interactions between immune and non-immune liver cells. Around the periphery of fibrotic regions, hepatocytes expressed many cholangiocyte-specific markers. While TREM2+ and monocyte-like macrophages were concentrated within fibrotic regions. In experimental validations, these macrophages exhibited a more suppressed and immunoregulatory phenotype. Thus, identifying multiple cell-cell interactions involving both pro-inflammatory and immune-regulatory cell-types that contribute to immune dysregulation in PSC.

## Methods

### Human Liver Tissue

Healthy human liver tissue from the caudate lobe was obtained from neurologically deceased donor livers acceptable for liver transplantation and without evidence of histopathological liver disease (neurologically deceased donor (NDD)). Samples were collected with institutional ethics approval from the University Health Network (REB# 14-7425-AE). PSC samples were collected from explanted tissue at the time of transplantation with institutional ethics approval from the University Health Network (REB# 20-5142). All patient clinical characteristics in **Supplementary Table 1**.

### NDD and PSC Sample Collection for scRNAseq & snRNAseq

Caudate lobes of 24 NDD livers and 4 PSC livers were collected and processed for single cell RNA sequencing (scRNAseq) fresh, as previously described(2,22) with a collagenase dissociation protocol(22). TLH was stored in FBS 10% DMSO in liquid nitrogen for further experimentation. Within 1 hour of caudate removal from the donor organ, intact caudate tissue sections were snap frozen and stored in liquid nitrogen, or embedded in optimal cutting temperature (OCT) medium to be stored at −80C(23). Nuclei for single nucleus RNA sequencing (snRNAseq) (3 PSC explants) were extracted from snap frozen tissue as described previously(3,24). All patient clinical characteristics in **Supplementary Table 1**.

### Preprocessing of each 10X Chromium sample

Sequencing reads were quantified using cellranger mapping to hg18 (for specific versions see **Supplementary Table 1**). Droplets containing viable cells were identified using EmptyDrops from the DropletUtils (v1.2.0) package(25). Since washing or sorting prior to loading cells into the 10X Chromium was not performed, identification of viable cells from cellular debris and ambient RNA was performed computationally. EmptyDrops were used with the following parameters: lower=20,000, niters=100,000, ignore=10, retain=100. Viable cells were defined using a 1% FDR threshold.

### Single-cell RNA sequencing QC

To ensure consistent quality across all samples, cells were further filtered with the following algorithm: cells with > 500 genes detected, > 875 total UMIs, and < 50% mitochondrial RNA were used for single-cell analysis. Cells from all samples were library-size normalized to 1500 UMIs/cell and log transformed using a pseudocount of 1. Each sample was scaled individually, and the top 2,000 most highly variable genes (HVGs) were identified in each sample using Seurat (v3.1.3)(26). Genes located in the mitochondrial genome were excluded from the set of HVGs and cells in each sample assigned to cell-cycle stages using Seurat’s default parameters. Cells were initially annotated using a custom algorithm (see supplementary methods) using our previous map(2).

### Data Integration

Raw, normalized, and scaled expression matrices for all samples were merged. Consensus highly variable genes were defined as those genes determined to be highly variable in at least two samples and that were detected in all samples. The samples were integrated using Harmony (v1.0)(27), with default parameters, based on the expression of the consensus highly variable genes. UMAPs were recalculated using the integrated lower dimensional space.

### Clustering and Annotation

The integrated NDD liver map was clustered using Seurat (v3.1.3)(26). The resolution parameter and k of the knn graph were varied from 0.3 to 2 and 40 to 80 respectively. The resulting clusterings were clustered using apcluster(v1.4.8)(28), with “p” = −2.5, using the variation of information(29) distance metric. This resulted in three exemplar clusters, which were compared using the uniformity of the automatic cell-type annotation of each sample to identify the optimal clustering: res = 0.6, k = 50. The resulting clusters were manually annotated by comparing markers of each cluster from the FindMarkers function in Seurat with known cell-type specific genes (**Supplementary Table 2 & 3**).

The integrated PSC scRNAseq data was clustered using the top 30 principal components, k=20, res=2 and the integrated PSC snRNAseq data was clustered using the top 30 PCs, k=20, res=2. The resolution was increased for PSC samples to separate neutrophils from other myeloid populations. Clusters that represented the same cell-types were merged during manual annotation (**Supplementary Table 4**).

### Subclustering

The integrated NDD liver map clusters were classified into 8 general classes (**Extended Data Fig. 1**). Cells in each of these classes were subset and subclustered by repeating the entire clustering and integration pipeline (**Extended Data Fig. 2 - 5**). The subclusters were manually annotated using marker genes identified using the Wilcoxon-rank-sum test.

### Pathway analysis

Pathway analysis was performed using fgsea (v1.8.0) with 5% FDR and 100,000 permutations using the MSigdb Hallmark pathways, MSigdb Immune pathways, and Reactome pathways. Only terms with between 15 and 1000 annotated genes were considered. Pathways significant at a 5% FDR were reported as significant. The Jaccard index was calculated between genes associated with each pair of enriched pathways and used to manually collapse redundant pathways from among the top 20 significantly enriched pathways from each annotation source.

### Macrophage-Endothelial cell-cell communication

We used scmap(30) to determine the specific identity of macrophages and endothelial cells comprising the macrophage-endothelial doublets. Doublets were independently mapped to the macrophage subpopulations and endothelial subpopulations using the most specific marker genes for each cell population as identified during manual annotation (**Supplementary Table 2**). Only doublets that could be reliably assigned (similarity > 0.6 & consistent between Pearson and Spearman correlations) to both a macrophage and endothelial subpopulation were counted to determine the number of doublets containing each possible pair of subtypes.

### PSC vs Healthy

To compare cell-type specific differential expression between PSC and Healthy single-cell and single-nucleus data, we calculated pseudobulk gene expression for each cell-type in each sample by summing the respective raw UMI counts. Pseudobulks were compared between Healthy and PSC using edgeR’s exactTest(31) using only genes passing QC thresholds in both PSC and Healthy data and cell-types that could be reciprocally matched between the two conditions based on overlapping marker genes (**Supplementary Table 4 & 6**). Differentially expressed genes were identified using FDR < 0.05 (**Extended Data Fig. 6**). Pathway enrichments were calculated as above, and cell-cell interactions were identified using ligand-receptor interactions obtained from the KEGG Cytokine pathway (hsa04060).

### In Silico Flow Sorting

To compare our single-cell RNAseq data to our flow cytometry data, we performed *in silico* gating of our single cell data using thresholds based on the non-immune fraction: PTPRC > 0.1, CD68 > 0.1 in normal tissue, CD68 > 0.6 in PSC. PSC scRNAseq was enriched in immune expression overall compared to healthy, thus higher thresholds for “on” for each gene were determined based on maximum expression seen in non-immune cell types.

### Nanostring GeoMx Digital Spatial Profiling Platform (DSP)

PSC and NDD liver sections were sliced from OCT-embedded sections and submitted to NanoString for staining with selected morphological markers: DAPI (nuclei), KRT (epithelial cells), CD45 (immune cells), and CD68 (macrophages) (**Extended Data Fig. 7**). For this experiment, we employed the GeoMx Cancer Transcriptome Atlas that contains over 1800 immune- and cancer-related targets. Expression data was deconvolved into cell-type composition using proprietary Nanostring software and the marker genes from our healthy liver map (**Supplementary Table 2**).

### Visium Experimental Protocol

Tissue was prepared for Visium Spatial transcriptomics as previously described(3). Briefly, liver tissue was embedded in OCT media, frozen, and cryosectioned with 16-um thickness at −10°C (cryostar NX70 HOMP). Sections were placed on a chilled Visium Tissue Optimization Slide (10x Genomics) and processed following the Visium Spatial Gene Expression User Guide. Tissue was permeabilized for 12 minutes, based on an initial optimizations trial. Libraries were prepared according to the Visium Spatial Gene Expression User Guide and samples sequenced on a NovaSeq 6000.

### Visium Computational Analysis

Visium spatial transcriptomic data was mapped to the Human genome (GRCh38-2020-A), demultiplexed and quantified according to UMIs using spaceranger(v1.1.0). Mitochondrial genome transcripts and genes with no more than 2 UMIs in any spot in the entire tissue slice were removed. Gene expression was normalized using sctransform, and data visualized using Seurat(v4.0.2)(26). Transcriptomic data was clustered using the standard Seurat pipeline: highly variable gene detection, PCA, k-nearest-neighbour detection, Louvain clustering, and wilcoxon-rank-sum test for marker genes, using default parameters. Note: spatially variable genes largely overlapped with highly variable genes but excluded many B-cell and red blood cell markers. Zonation scores were calculated by first rotating principal components using the base R varimax function to improve their interpretability (See Supplementary Methods). Significance of zonation was calculated using linear regression between zonation score and the normalized gene expression. Colocalization was determined using Pearson correlation.

### Flow Cytometry & Cytokine Analysis

Cell suspension from frozen total liver homogenate (TLH) were stained as previously described(2,32,33) with live/dead Zombie Violet (Biolegend, 423114) or Zombie NIR (Biolegend, 423106) to assess viable cells and fluorophore-conjugated monoclonal antibodies to the following human cell-surface markers: anti-CD45-BV650 (Biolegend, 304044), anti-CD206-AF647 (Biolegend, 321116), anti-HLADR-AF700 (Biolegend, 307626), anti-CD68-R780 (Biolegend, 333816), and anti-CD68-PeCyp5.5 (Biolegend, 333814). Intracellular cytokine staining was performed to examine the functional differences in CD68^+^ cells that were either CD206^+^ or CD206^−^ in PSC and NDD TLH, cell suspensions (2×10^6^ cells) from the NPC fraction were stimulated in 12-well plates with 10 ng/ml LPS, 25ng/ml IFNg for 6 hours in the presence of BFA/monensin and intracellular secretion of TNFα, detected with anti-TNFα-PeCy7 (Biolegend, 502930) and anti-TNFα-PacBlue (Biolegend, 502920), was examined as previously described(2,32). Gating strategy for cell-surface markers was set based on background auto-fluorescence measured in unstained and fluorescence minus one (FMO) controls (**Extended Data Fig. 8, 9**), and gating strategy for intracellular markers in stimulated samples was set based on unstimulated controls and FMOs (**Extended Data Fig. 8, 9**).

### Histopathology of PSC samples

Tissue sections were formalin-fixed and paraffin-embedded for histological and immunohistochemical staining. Disease stage according to fibrosis was defined by the Nakanuma score and stage(34) by assessing the following components, 1) fibrosis, and 2) bile duct loss. Annotation of fibrotic areas and chronic biliary disease, referred to as scars in this study, within the liver parenchyma of PSC explant livers was defined based on Masson’s Trichrome stain (highlighting fibrosis) and Cytokeratin 7 (CK7) immunohistochemical stain (highlighting biliary metaplasia in hepatocytes as a feature of chronic biliary disease) (**Extended Data Fig. 10**)(35). CK7 staining highlights the hepatocytes with biliary metaplasia near the edges of fibrotic areas (scars) in PSC explant samples (**Extended Data Fig. 11**).

## Results

### Global Healthy Map

To generate an expanded reference single-cell liver atlas suitable for identifying PSC-aberrant transcriptomic patterns, we collected single-cell transcriptomes from over 100k single-cells from 24 different healthy livers with equal representation of males and females and spanning a wide age range (**Figure 1**). Cells were dissociated as previously described(2) from the caudate lobe of NDD donors prior to organ transplantation to a living recipient. ScRNAseq was generated using the 10X Chromium with both 3’ and 5’ chemistry (**Supplementary Table 1**, **Figure 1 C**). Samples integrated into a single atlas, which was clustered into 20 coarse-level cell groups. We found all samples merged and contributed to almost every cell cluster in our map and cellular phenotypes were remarkably consistent across demographic factors (Pearson correlation between donors > 0.5 in 17/20 clusters) (**Figure 1 G**, **Supplementary Table 2 & 3**). Using compositional analysis (see Methods), we found significant variations in cell-type frequencies across samples (**Figure 1 F**). However, this is likely due to dissociation and tissue sampling effects rather than donor-specific differences, as significant associations between cell-type frequency and donor sex or age were not identified (linear regression Benjamini-Hochberg adjusted p-value > 0.3). These coarse level cell-types were subclustered into a final total of 38 cell-states (**Extended Data Fig. 2-3**). These included tissue-resident and circulating NK cells, naive and plasma B-cells, and an ultra-rare population of mucus-producing cholangiocytes.

**Figure 1:**
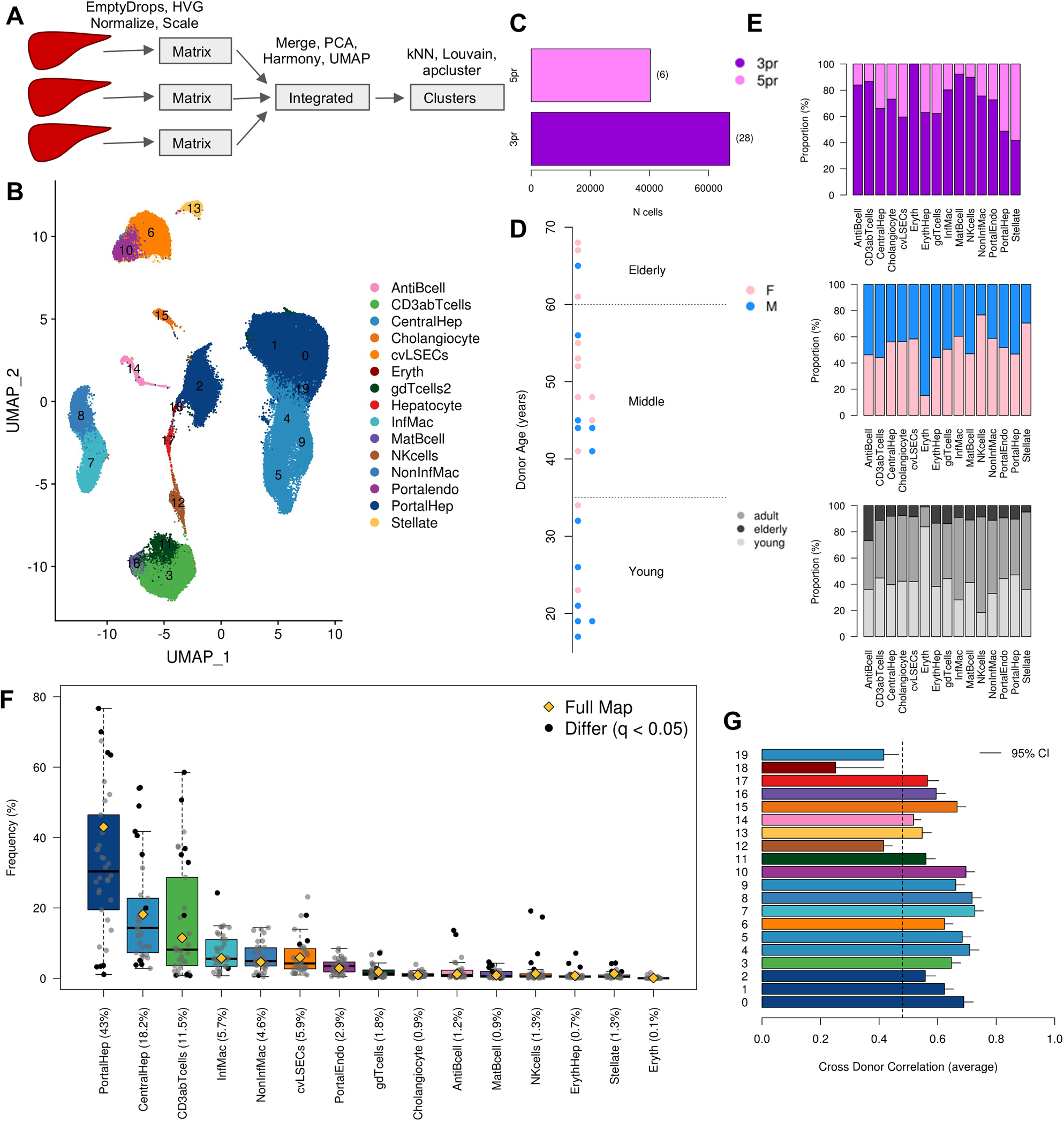
100k single-cell map of Healthy Human Liver reveals conserved cell-types across donors. (A) Single-cell RNAseq sample collection and analysis workflow (B) Integrated UMAP, containing 20 clusters annotated to 15 major cell-types (C) Number of cells sequenced with 5’ scRNAseq vs 3’ scRNAseq, numbers in brackets indicate the number of samples sequenced with each technology. (D) Age and sex profiles of the 24 donor livers pink = female, blue = male. ((E) Proportion of cells from different technologies, sexes and ages contributing to each cell-type of the integrated data. Black points indicate significant enrichment / depletion (p < 0.05) based on a hypergeometric test using compositional analysis. There were no significant associations between demographic characteristics and the frequency of different cell-types. (G) Average correlation of the cluster profiles between different donors. The cluster profile is the mean expression of cells within that cluster. Dashed line indicates the average correlation between the cluster profiles of different clusters within the same donor.

### Expanded healthy liver map elucidates human macrophage diversity

Our original liver atlas was only able to distinguish two different macrophage populations characterized by opposing relationships with inflammation(2). However, several more distinct macrophage phenotypes were evident after subclustering the 11,127 macrophages of our expanded map (**Figure 2 A**). We identified 15 subclusters among the macrophages in our map, several of these exhibited simply different expression in inflammatory (*S100A8, S100A9, S100A6, VCAN, LYZ*) or known non-inflammatory (*CD5L, MARCO, VCAM1*) markers. These clusters were merged together due to their lack of unique marker genes, leaving 10 distinct phenotypes of macrophages that were conserved across donors and that were consistent with the macrophage subtypes identified in Guilliams et al.(4) (**Extended Data Fig. 4**). After merging, non-inflammatory Kupffer cells and inflammatory monocyte-derived macrophages accounted for 40% of all macrophages.

**Figure 2:**
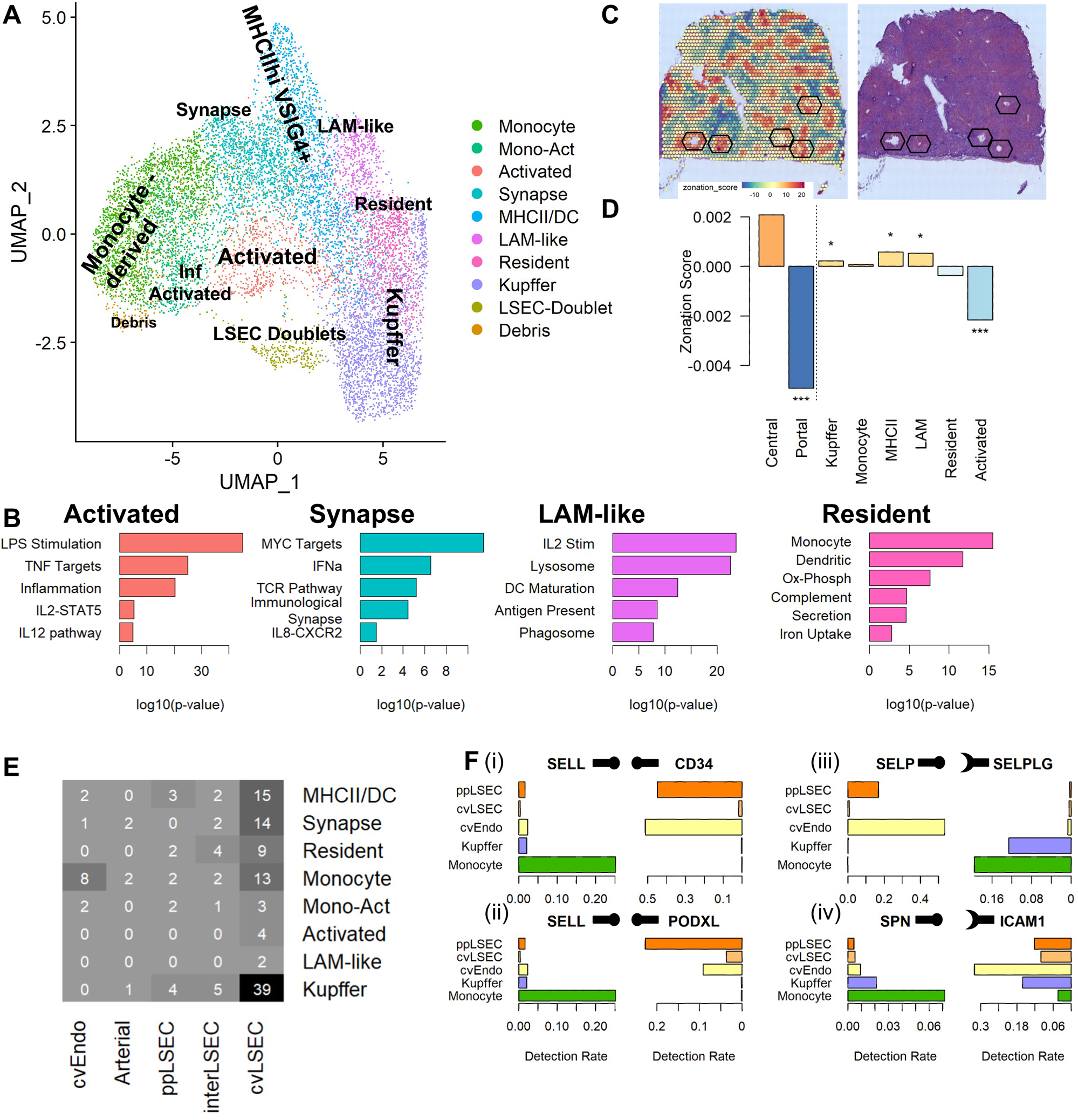
Macrophage diversity revealed by expanded healthy Liver map. (A) 11,127 macrophages were subclustered using Seurat and manually annotated to identify 10 distinct phenotypes. (B) Each subtype was enriched for distinct functional pathways as determined by GSEA. (C) Varimax PCA identifies a component that quantifies the zonation of each spot. Black hexagons indicate pericentral veins of distinct lobules. (D) Using the zonation score across four slices of spatial transcriptomics we determine non-inflammatory macrophage populations are pericentral while activated macrophages are periportal. (E) Number of doublets assigned to each specific pair of subtypes using scmap. (F) Most specific ligand-receptor interactions from Cellphonedb for Monocyte-derived and cvEndo cells, nobs indicates proteins labelled as ligands and cups indicate proteins labelled as receptors.

The remaining 60% were spread across 8 rarer phenotypes, including LYVE1^+^ FOLR2^+^ TIMD4^+^ non-inflammatory resident macrophages that were similar to the self-renewing resident cardiac macrophages found in mice(36,37) and MHCII-high VSIG4^+^ macrophages that were similar to those observed in many mouse tissues(38) (**Figure 2**). Among the MHCII-high population, we identified 10 cells that were XCR1^+^ CLEC9A+ which may represent cDC1 / Cross-presenting dendritic cells(39) (**Extended Data Fig. 4 C**). Whereas CLEC10A showed low expression among the other MHCII cells suggesting they may represent cDC2s.

In addition, we identified a novel macrophage subtype characterized by the expression of *PLAC8, LST1, IFITM3, AIF1, COTL1* in addition to some inflammatory markers (*FCN1, LYZ, S100A4, S100A8*). *COTL1* and *LST1* are cytoskeleton-related proteins involved in lamellipodia at immune synapses(40,41). *CD52*, another top marker of this cluster, is involved in interactions with T-cells(42). Pathway enrichment analysis revealed upregulation of interferon-alpha (IFNa) response, the TCR signaling pathway, the immunological synapse and *IL8-CXCR2* signaling (**Figure 2 B**). These results suggest these inflammatory macrophages are specialized in interacting with and activating T cells.

The rarest subtype of macrophages (5%) uniquely expressed *GPNMB, TREM2, FABP5, ACP5, PLD3 and LGMN* similar to previously described ‘lipid-associated’ macrophages (LAM-like)(4,43). However, we found these macrophages lacked expression of some LAM-associated markers: *CD93, MS4A7, and FCGR2B* (**Figure 2**), and were not enriched in lipid-associated pathways. Instead, pathway analysis found an enrichment of phagosome and lysosome related transcripts including the lysosomal proteins *CP5, PLD3, PSAP and CSTB*, suggesting these macrophages are specialized in phagocytosis(44) (**Figure 2 B**).

In addition to these subtypes of macrophages, we observed a type-independent activation pathway in macrophages. This signature was characterized by high expression of *IL1B, CD83, CXCL2, CXCL3, NAMPT, THBS1,* and *AREG*, as well as genes involved in pathways associated with in vitro LPS stimulation of macrophages and TNFα signaling (**Figure 2 B**). This pathway was observed in both monocyte-like macrophages and macrophages lacking any other phenotype-signature.

### Macrophage-endothelial cell-cell communication

Previous work has shown that liver sinusoidal endothelial cells (LSECs) are important in recruiting monocytes to the liver after injury, and LSECs contribute to inducing a Kupffer cell-like phenotype in recruited monocytes if native Kupffer cells are artificially depleted in mouse(45,46). Interestingly, we found a significant number of doublets between macrophages and LSECs in our healthy human liver that expressed macrophage markers (*CD163, CD68, TIMP1,* and *C1q*) and LSEC markers (*DNASE1L3, ENG, SPARC, CLEC1B*). These cells accounted for 282 cells (3% of all macrophages), which was 10-fold higher than we expected based on the loading of our samples and frequency of macrophages and LSECs in our map (expect: 0.3%, p < 10^-100^). Using scmap(30), 142/282 doublets were reliably assigned to both a specific macrophage subcluster and a specific endothelial subcluster (**Figure 2 E**). Of these the most common pairwise interaction was between Kupffer cells and central venous LSECs (cvLSECs). However, Monocyte-like macrophages were uniquely likely to form doublets with central venous endothelial cells (cvEndo) suggesting these represent newly recruited monocytes to the liver.

To confirm these doublets represent cell-cell interactions we used CellPhoneDB(47) Results were filtered to identify interacting proteins involved in cell-cell adherence or close-contact cell-cell communication that were specific to cvEndo and Monocyte-like macrophages (**Figure 2 F)**. The 10 top interactions after filtering included *SELL-PODXL*, which are known to be involved in immune cell rolling-adhesion(48), *ITGAL/SPN-ICAM1* that have previously been shown to be important for tissue infiltration of immune cells(49), and *SELP-SELPG*, a known mediator of leukocyte recruitment(50) (**Figure 2**). These results indicate Monocyte-like macrophages represent monocytes, recently recruited to the liver through the central vein.

### Spatial transcriptomics identifies spatial location of macrophage populations

To add spatial resolution to our single-cell atlas, we sequenced four consecutive slices using VISIUM spatial transcriptomics (**Figure 2 C,D**, **Extended Data Fig. 5**). Hepatocyte zonation was captured as either the first or second principal component in each slice. This score was used as weights to calculate the average zonation of individual genes which were summed to determine the average enrichment of specific cell-types. These results confirmed our endothelial cell annotations, and indicated NK and T cells and B cells were weakly periportal (**Extended Data Fig. 5 E-F**). Examining macrophage subsets, we found MHCII^+^, LAM-like macrophages and Kupffer cells were generally pericentral, while activated macrophages were periportal (**Figure 2 D**).

### Transitioning hepatocytes are present around late-stage PSC lesions

We sequenced 12,000 single cells using 5’ scRNAseq from 4 PSC livers and 23,000 nuclei from 3 PSC livers (**Figure 3 A, Figure 4 A**). These were integrated into separate single-cell and single-nucleus maps using the same pipeline as our healthy map in **Figure 1**. In agreement with our previous findings (3), snRNAseq captured a more representative sample of cells present in diseased liver with 64% of nuclei originating from hepatocytes, whereas hepatocytes made up only 10% of the scRNAseq map. Of these hepatocytes we observed a decrease in the frequency of periportal hepatocytes in PSC livers relative to the healthy livers (28% vs 49%, Fisher’s exact test: p < 10^-300^), but only modest differences in proliferation (**Figure 3 B**). In addition to pericentral and periportal-like hepatocytes, we identified a novel hepatocyte phenotype only found in our PSC samples (Hepato2). The markers of this hepatocyte cluster had a significant overlap with markers of the cholangiocyte cluster: *BICC1, DCDC2, FGF13, THSD4, RASEF, GALNT18, ENAH, LAMA3, C14orf105, BIRC3, CXCL8, CREB5* (hypergeometric test, p < 10^-10), suggesting these may be hepatocytes transforming into a cholangiocyte-like phenotype. These hepatocytes also expressed many unique markers including: *SQSTM1, AKR1B10, ANGPTL8, KRT23, PLA2G4C, AFF3, DAPK2, SULT1C2, HKDC1* (**Figure 3 C**). We confirmed the presence of transitioning hepatocytes around the periphery of areas of PSC-related onion skin fibrosis, as annotated by a qualified pathologist, with high-resolution imaging of *KRT7* (CK7) (**Extended Data Fig. 10, 11**), which we refer to as scars using VISIUM spatial transcriptomics. Interestingly, these hepatocytes were only present around scars lacking any surviving cholangiocytes and which had a rough, nuclei-rich edge when examined histologically (**Figure 3 D-H**).

**Figure 3:**
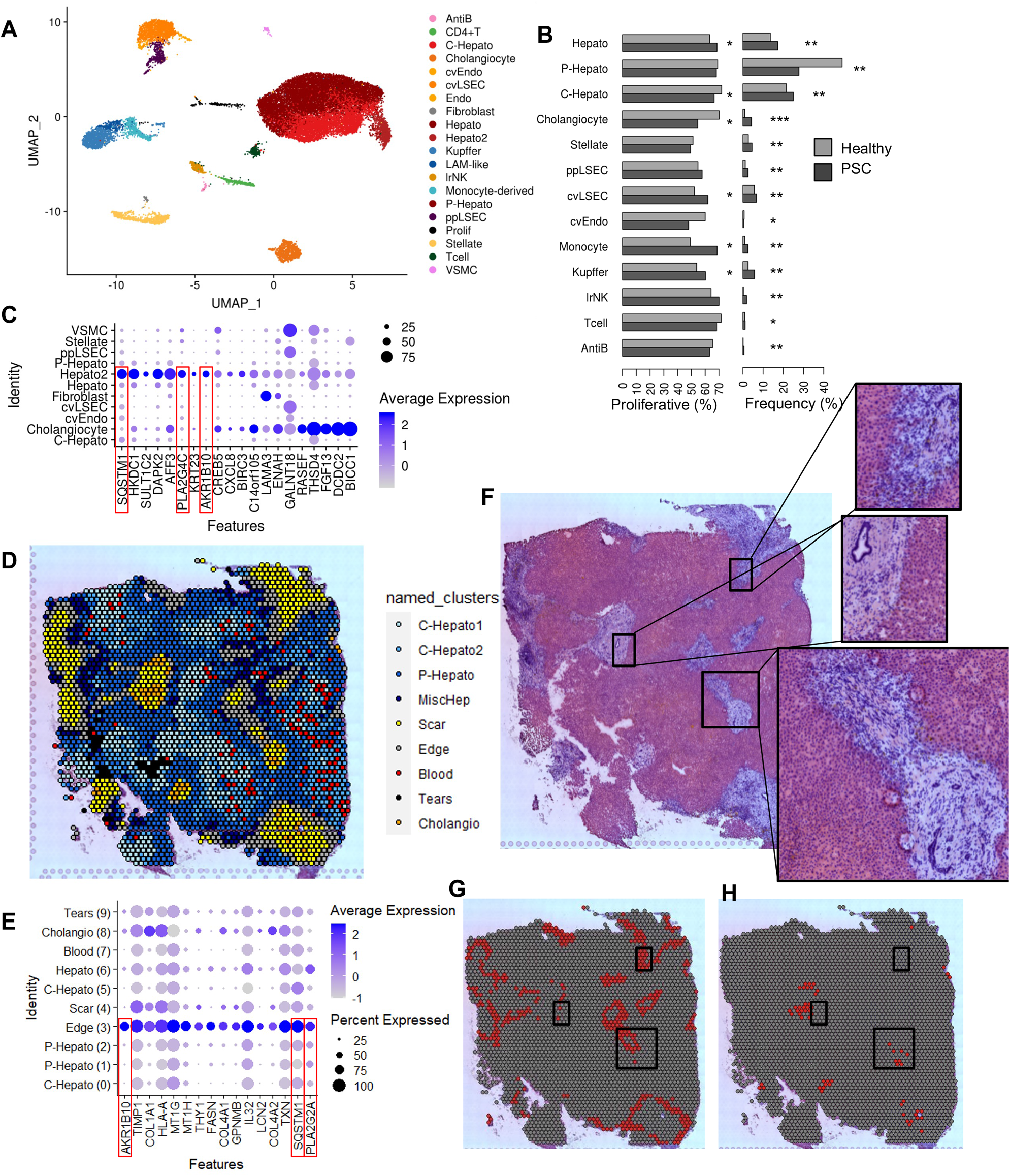
Edges of PSC scars harbour unique transitioning hepatocytes. (A) Integrated map of 23,000 nuclei from 3 livers with PSC. (B) Proportion of proliferating cells (left) or proportion of the map (right) for each cell-type found in both the PSC and Healthy snRNAseq maps. Significance obtained from proportion test: * p value < 0.05, ** p value < 10^-10^, ***p value < 10^-100^(C) Top marker genes of the unique PSC hepatocytes (Hepato2), red rectangles indicate markers shared with the scar edges in panel E. (D) Clustering of PSC VISIUM spatial transcriptomics data. (E) Top markers of scar edges, red rectangles indicate markers shared with PSC specific hepatocytes. (F) H&E stained image of the PSC tissue slice used for spatial transcriptomics. Squares zoom into various scars with and without surviving bile ducts. Scars with bile duct exhibit a smooth edge while those without exhibit rough nuclei-rich edges. (G & H) Highlight of spatial transcriptomic clusters: Scar edges (G) and Cholangiocytes(H), rectangles indicate zoom in regions seen in F, note that only rough edges are included in the “Edge” cluster.

**Figure 4:**
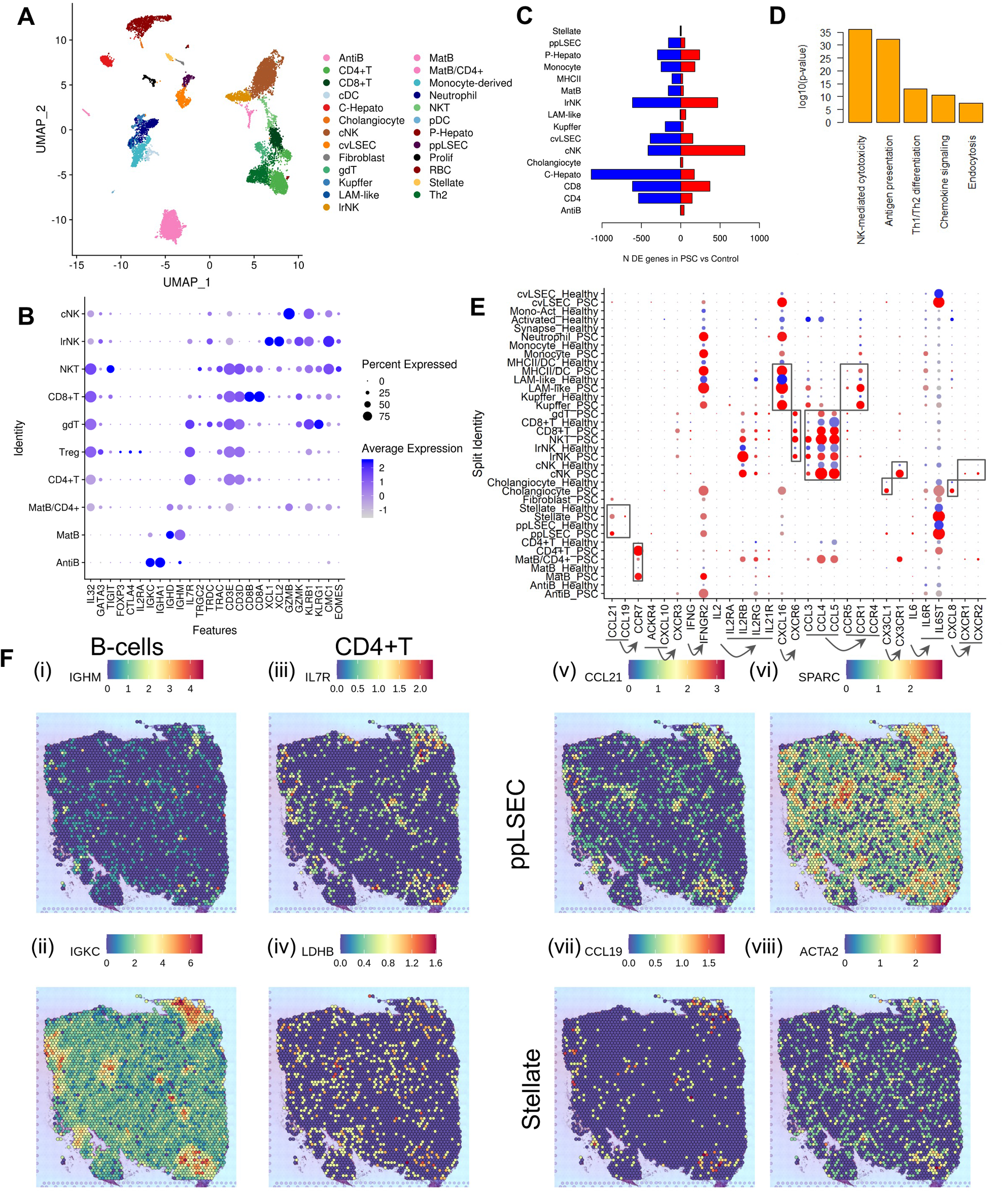
Single-cell RNAseq reveals immunological complexity of PSC. (A) Integrated map of 12,000 cells from 4 PSC livers. (B) Marker genes of PSC-specific lymphocyte populations. (C) Number of differentially expressed genes between healthy and PSC cells, according to edgeR applied to cluster-specific pseudobulks (5% FDR), red are genes up regulated in PSC, blue are genes down regulated in PSC. (D) Top pathway enrichments for genes upregulated in cNK cells from GSEA. (E) Expression of ligand-receptor pairs for known cytokines from the KEGG database in PSC and healthy single cell maps. Arrows indicate ligand to receptor relationships. (F) VISIUM spatial transcriptomics of key marker genes of immune populations. (i) naive B cells (ii) plasma B cells (iii-iv) CD4+T (v-vi) ppLSECs (vii-viii) hepatic stellate cells.

### Immune expansion within PSC scar regions

While hepatocytes, stellate cells and cholangiocytes were well represented in the single-nucleus map, lymphocyte populations were difficult to distinguish. In contrast, our single-cell map revealed the presence of both major types of dendritic cells and a large diversity of lymphocyte populations, including many that were not present in our healthy map (**Figure 4 A,B**). Disease-specific lymphocyte populations included NKT-like cells that expressed lrNK markers (*EOMES, CMC1, KLRG1, KLRB1*)(51), cNK markers (*GZMK, XCL2*)(52) and CD8^+^ T cell markers (*CD8A, CD8B, CD3D,TRAC, TRGC2*). DoubletFinder(3) determined this population was not doublets (estimated 3% doublets). A population of CD4^+^ T regulatory cells (*IL2RA, CTLA4, FOXP3, TIGIT, GATA3*)(53), these cells expressed IL32 which has been previously linked to nonalcoholic fatty liver disease(54) and the expression of FOXP3 by Tregs in cancer(55). We found a population of gamma delta T cells (*CD3E, TRDC, TRGC2, TRDV2, TRGV9*) similar to the NK T cells which expressed a combination of T cell (*IL7R, CD3D, TRAC*) and NK cell (*KLRB1, KLRG1*) markers.

We also identified a cluster of CD4^+^T cell and mature naive B cell doublets (MatB/CD4^+^, estimated 91% doublets). These naive B cells expressed more IGHD whereas the naive B cells, not present in these doublets, expressed more IGHM. This is intriguing as recent work has shown that IGHD^+^ B cells react less strongly to “self” antigens than IGHM^+^ B cells in mice(56). Spatial transcriptomics confirmed co-localization of CD4^+^ T cells and B-cells in fibrotic regions (**Figure 4 F**). Differential expression analysis between 4 healthy 5’ single-cell samples from our integrated map suggested a rapid expansion of antibody-secreted B cells (AntiB) with 8 out of the 10 most upregulated genes in PSC being specific B cell receptor (BCR) variable segments (**Supplementary Table 4**).

To examine inflammation related cell-cell signaling, we examined the expression of gene pairs KEGG cytokine pathways (hsa04060). We found evidence of interactions between CCR7^+^ CD4^+^ T cells and CCL19^+^ Stellate or CCL21^+^ ppLSEC cells in fibrotic regions (**Figure 4 E, F**). The colocalization of *CCL19, CCL21* and *IL7R* (CD4^+^ T cells) was confirmed in our VISIUM spatial transcriptomics (Pearson correlation 0.2, p < 10^-30, **Figure 4 V-VII**), and across multiple replicates of Nanostring (**Figure 5 C-D**). Using Nanostring we observed the colocalization of CD4^+^ T cells, Stellate cells, ppLSECs and Mature B cells in PSC. However, this was only the case for the advanced diseased regions examined, and large-scale heterogeneity was observed between different regions of interest from the same original liver samples.

**Figure 5:**
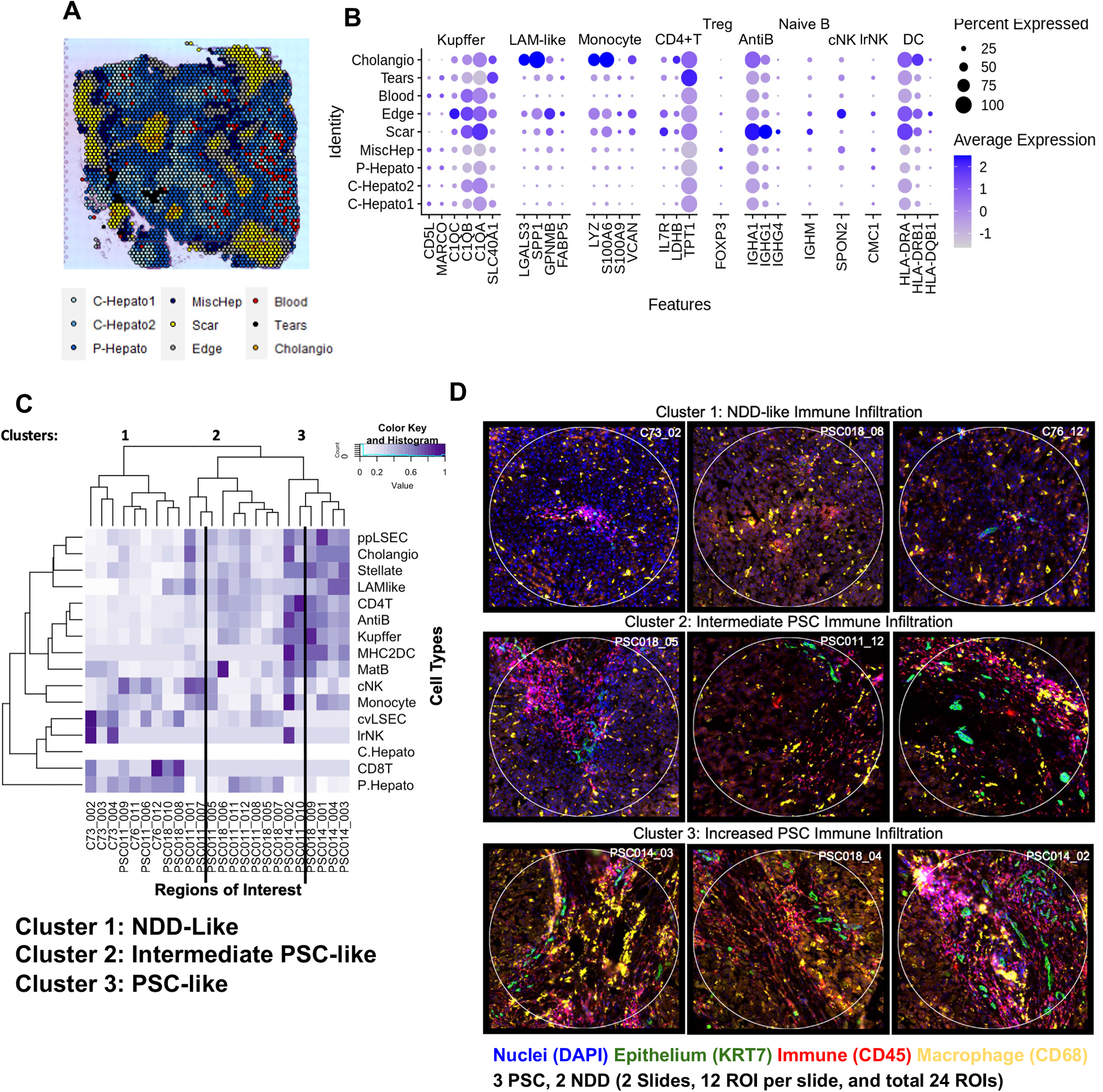
Enrichment of Immune cells in PSC (fibrotic areas) (A) VISIUM spatial transcriptomics was clustered identify layers of distinct regions around fibrotic areas in PSC. (B) Expression of marker genes for each lymphocyte and macrophage subtype from the PSC and healthy single-cell maps in the PSC VISIUM data. LAMs and Monocytes were enriched in the core of scars near cholangiocytes and periphery of fibrotic areas, while B cells, T cells, and antigen presenting cells were enriched in the center of fibrotic regions, whereas NK cells present at the periphery. (C) Deconvolution of Nanostring digital spatial expression using cell-type markers from the healthy and PSC single-cell maps. (D) Images of typical example regions for each cluster identified in Nanostring. Nuclei are shown in blue, cholangiocyte epithelium in green, leukocytes in red, and tissue resident macrophages in yellow. Each circle is 660um in diameter. Complete NanoString ROIs are in Supplementary Figure

However we found the highest amount of differential expression between PSC and healthy liver within the NK populations, and in particular cNK cells (**Figure 4 C,D**). Pathway analysis revealed these genes were involved in NK cell cytotoxicity, and antigen presentation. Among these genes upregulated in cNK cells we identified CXCR1/2 and CX3CR1 chemokine receptors (CXCR2 l2fc = 2.3, CX3CR1 l2fc = 2.4, FDR < 5%), the ligand of which were specifically up regulated in PSC Cholangiocytes (CXCL8 l2fc = 3.12, FDR < 5%, CX3CL1 = 0.55, FDR > 5%). However, since both ligands are soluble secreted factors, spatial transcriptomics could not confirm this putative interaction.

### Macrophage dysfunction in PSC livers

Unlike the lymphocyte populations which upregulated inflammatory pathways in PSC, macrophages increased inhibitory signaling molecules in PSC, including *IL27RA, VSIR,* and *IL18BP*. *IL27RA* has been shown to suppress cytokine production in macrophages(57,58) and was upregulated in PSC Kupffer cells. *VSIR* was upregulated in Kupffer cells and LAM-like macrophages, and is known to reduce the responsiveness of macrophages to LPS stimulation (59). Monocyte-like macrophages adopted a fibrosis-associated phenotype upregulating *LGALS3, TREM2*(*60*), *SPP1*(*7*) and *ADA2*(*61*). We confirm this scar-associated phenotype of monocyte-like macrophages using our spatial transcriptomics which indicated enrichment of these cells around cholangiocytes in the center of regions of fibrosis including concentric periductal fibrosis (**Figure 5**). In addition we observe that TGFβ, a potent suppressor of macrophage function(62), co-localized with these cells within those regions (**Figure 5**).

Comparing Kupffer cells in our snRNAseq and scRNAseq, we identified three genes significantly upregulated in both assays all of which are involved in phagocytosis and possibly macropinocytosis: *SRGAP1*(*63*), *SRGAP2*(*64*), *APBB1IP*(*65*). For the Monocyte population, 15 genes (**Supplementary Table 5**) were consistently upregulated in PSC in both single-cell and single-nucleus maps, several of which have been previously reported as upregulated in fibrosis (*PLA2G7, LGMN, HSPH1)* or suppression of inflammation (*ZBTB16, SNX9*). In addition, we found *AIF1* significantly upregulated in the monocyte-like myeloid population in PSC, which is consistent with recent reports of bulk RNAseq profiling of PSC explants(66).

Flow cytometry of primary patient macrophages revealed a significant depletion of CD68^+^ cells relative to CD45^+^ (PTPRC) cells in PSC which was confirmed with *in silico* gating of our single-cell data (**Figure 6 A-C**). Additionally, we observed increased percentages of CD206^+^ macrophages in PSC in comparison to NDD (**Figure 6 D-E**), as previously described in PSC(20). Intracellular cytokine staining (ICS) stimulation assays indicated a significantly reduced capacity of PSC macrophages to secrete TNFα in response to LPS and IFNγ stimulation (**Figure 6 F-G**). This is consistent with the upregulation of inhibitory signals seen in our single-cell data, and the colocalization of TGFβ with monocyte-derived macrophages in our VISIUM data (**Extended Data Fig. 8, 9**). Another possible mechanism of this suppressed macrophage phenotype is direct signaling by contact between macrophages and bile acid(67).

**Figure 6.**
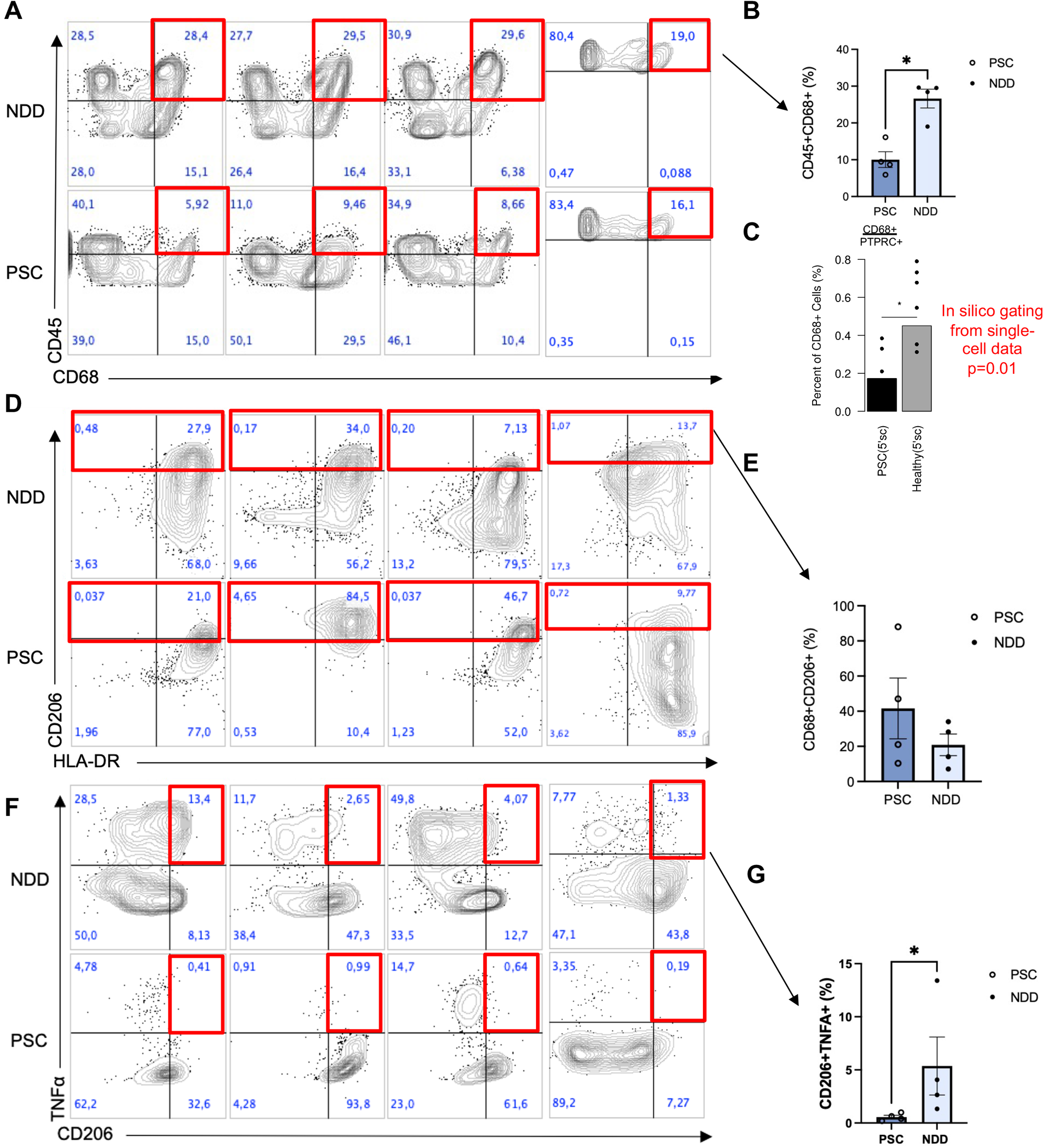
PSC macrophages are dysfunctional in cytokine secretion. A) Flow gating of CD45/PTPRC+ and CD68+ in PSC and NDD TLH. B) PSC samples contained significantly fewer double positive cells (p < 0.05). Percentage of intact flow sorted cells that are CD68+CD45+ in the PSC and NDD samples. C) *In silico* gating of scRNAseq replicates decrease in CD68+CD45/PTPRC+ cells (p=0.01). D & E) Flow gating of CD45+CD68+ cells reveals increase in CD206+ subset in PSC (P>0.05). F & G) Following LPS stimulation, PSC macrophages (CD45+CD68+) exhibit reduced production of TNFa (p < 0.05). Statistical significance for flow cytometry and intracellular cytokine staining is evaluated using a non-parametric Mann-Whitney U Test, ∗∗∗P < 0.001, ∗∗P < 0.01, ∗P < 0.05, error bars represent the standard error of the mean (SEM). Full gating strategy and unstimulated controls are shown in **Supplementary Figure 6**.

## Discussion

The complex cellular landscape of the PSC liver is relatively poorly understood but is being better elucidated through tissue mapping with single-cell technologies(11,13–15). In our investigation, we leverage a 24-sample NDD liver map (**Figure 1**) and assess end-stage PSC through single-cell, single-nucleus, and spatial transcriptomics to delineate the cellular pathways of liver injury in PSC. We uncovered 38 distinct cell-types and cell states in the NDD liver, including a pericentrally located myeloid population similar to the previously described lipid-associated macrophages(4,43), as well as myeloid subpopulations associated with immunological synapses and a consistent activation signature (**Figure 2**). In PSC livers, we identified a further 8 disease-associated cellular states including NKT-like cells, neutrophils, dendritic cells, and a subpopulation of hepatocytes (**Figure 3-4, Supplementary Table 6**).

These immune populations are likely a result of *in situ* expansion and recruitment from the blood into the diseased liver tissue (12,13,15). As previously reported for healthy liver tissue, immune subpopulations could only be identified in scRNAseq (**Supplementary Table 7**), whereas the hepatocyte subtype was only detected in snRNAseq (**Supplementary Table 8**)(3). Many of the immune populations were colocalized within fibrotic ‘scar’ regions in PSC, including CD4^+^ T cells and antibody secreting B cells, this is consistent with bulk RNAseq findings of enriched B lymphocytes and FOXP3^+^CD4^+^ T cells in the PSC liver(66)

In contrast, we observed disease-associated hepatocytes were specifically located in the border regions surrounding the fibrotic ‘scar’ regions in PSC (**Figure 3**). These hepatocytes expressed many markers associated with cholangiocytes, suggesting they may be transitioning to cholangiocytes. Previous studies have demonstrated hepatocytes undergo this transition in the absence of functional bile ducts in a TGFβ-mediated manner(68,69). Consistent with these results, TGFβ was expressed in the PSC fibrotic regions, particularly in scars with transitioning hepatocyte edges (**Extended Data Fig. 12**).

TGFβ has also been implicated in the suppression of macrophage activation(70) and the progression of cholestasis(71,72). Macrophages colocalized with TGFβ in our PSC spatial transcriptomics suggesting possibly suppression. We validated this prediction experimentally and observed suppressed TNFα secretion cytokine secretion and inflammatory potential by CD206^+^ KC-like myeloid cells in PSC following stimulation (**Figure 6**). A similar cytokine secretion dysfunction has been reported in inflammatory bowel disease (IBD), a common comorbidity of PSC(73,74). This suggests a possible mechanism by which macrophage dysfunction may underlie PSC and IBD pathogenesis(13,16,20).

Like most studies on PSC(11,12,18), our data was mainly limited to a single region of each end-stage explanted liver, profiled across multiple modalities, which was compared to non-diseased livers. However, our Nanostring spatial transcriptomic data, which sampled multiple regions of the same liver, allowed us to uncover substantial heterogeneity in disease phenotype across even relatively small regions of these livers (**Figure 5**). PSC is generally known to be a slow progressive disease; thus, it remains to be determined how and when the cell signatures and cellular dysfunction we describe in this work develop in the trajectory of disease and whether these patterns are specific to PSC or general features of late-stage cholestatic liver disease. Thus, future work will expand this map to recover temporal and spatial heterogeneity of disease. Our results demonstrate the power of combining multiple high-resolution gene expression platforms including 3’ single cell, 5’ single cell, single nucleus, and spatial transcriptomics to fully elucidate the interactions between tissue microenvironment, cellular identity, and immunological function underlying a complex inflammatory disease.

## Abbreviations

FBS: fetal bovine serum
DMSO: dimethyl sulfoxide
FMO: fluorescence minus one
HVG: highly variable genes
FDR: false discovery rate
AntiB: antibody secreting B cells, also known as Plasma cells, that express CD79A, CD79B and various light and heavy chains of the B cell receptor (probably as soluble antibodies) e.g. IGKC, IGHG, IGLC, IGHA1/2
BCR: B cell receptor
CD45: cluster of differentiation antigen 45, originally called leukocyte common antigen, the protein produced by the PTPRC gene in humans.
CD206: the protein produced by the MRC1 gene, a marker of tissue resident macrophages
cvEndo: central venous endothelial cells
CK7: the protein produced by KRT7 gene.
ICS: Intracellular cytokine staining
ICU: Intensive care unit
LAM: lipid associated macrophage
LPS: Lipopolysaccharides
LSEC: liver sinusoidal endothelial cells
ppLSEC: periportal liver sinusoidal endothelial cells
cvLSEC: central venous liver sinusoidal endothelial cells
IBD: inflammatory bowel disease
IFNγ: Interferon gamma
MHCII: MHC Class II molecules, a class of major histocompatibility complex (MHC), includes HLA-DRA, HLA-DPA1, HLA-DQB1, HLA-DPB1.
NDD: neurological determination of death
NK cells: natural killer cell
cNK: circulating natural killer cells
lrNK: liver resident natural killer cells
NKT: natural killer T cells
PSC: Primary Sclerosing Cholangitis
scRNAseq: single-cell RNA sequencing
snRNAseq: single-nucleus RNA sequencing
ST: spatial transcriptomics
TLH: Total liver homogenate
TNFα: Tumor necrosis factor alpha
TBFβ: Transforming growth factor beta
UMI: unique molecular identifier
All other abbreviations are official HGNC gene symbols.

## Data Availability

The healthy liver map data and the spatial transcriptomic data are available for exploration with our interaction shiny tools (see links below). Seurat objects and raw count matrices for all of these data and the PSC single-cell and single-nucleus data have been uploaded to cellxgene and will be publicly available (currently pending manual data curation). Raw reads for all data is in the process of being uploaded to GEO. Accessions for all these data will be included in the final published version of this manuscript.

## Healthy Map Interactive Browsers

https://macparlandlab.shinyapps.io/generalmap/

https://macparlandlab.shinyapps.io/macrophages/

https://macparlandlab.shinyapps.io/lymphocytes/

https://macparlandlab.shinyapps.io/endothelial/

https://macparlandlab.shinyapps.io/cholangiocytes/

https://macparlandlab.shinyapps.io/b_cells/

https://macparlandlab.shinyapps.io/stellate/

## Spatial Healthy

https://macparlandlab.shinyapps.io/healthylivermapspatialgui/

## Financial Support Statement

Funding provided in part through grant number CZF2019-002429 from the Chan Zuckerberg Initiative DAF, an advised fund of Silicon Valley Community Foundation; an early researcher award from the Government of Ontario (ERA19-15-210); funds from the University of Toronto’s Medicine by Design initiative, which receives funding from the Canada First Research Excellence Fund (CFREF) to SAM, GDB and IDM; by the NRNB (U.S. National Institutes of Health, grant P41 GM103504) to GDB and by the Toronto General and Western Hospital Foundation. DN has received graduate fellowships from CIHR (CGS-D). CTP has received postdoctoral funds from the Canadian Network on Hepatitis C (CanHepC). CanHepC is funded by a joint initiative of the Canadian Institutes of Health Research (CIHR) (NHC-42832) and the Public Health Agency of Canada (PHAC). CTP received postdoctoral funds from PSC Partners Seeking a Cure Canada which was matched by the UHN foundation. Aspects of this work were performed in collaboration with the Canadian Autoimmunity Standardization Core (CIHR-HUI-159423). This work was funded in part by CIHR grants PJT162298 (IDM) and PJT162098 and HIT168002 (SAM). SAM holds a CRC Tier 2 in Liver Immunobiology.

This publication is part of the Human Cell Atlas –www.humancellatlas.org/publications/. The authors acknowledge the Princess Margaret Genomics Centre for support and services.

## Author Contributions

S.A.M., G.D.B. and I.D.M designed the study. T.S.L, D.N, X.-Z.M., B.K.G., L.L, conducted experiments. E.W., B.A., S.M., A.R., and G.H. recruited and consented patients. E.W. S.M., B.S., G.H.M. and I.D.M harvested the surgical specimen. D.N., C.T.P., L.L., D.C. and S.C, X.-Z.M., and J.M. expedited the processing of the harvested surgical specimens for storage and experiments. C.T. interpreted the histology. T.S.A., D.N., L.L., S.C., D.C., B.H., A.G., A.G., A.R., G.M.H., S.A.M., G.D.B., and I.D.M. analyzed and interpreted the data. T.S.A, D.N., S.A.M., G.D.B. and I.D.M prepared the manuscript with critical revision from all authors.

## Supplementary Figures

**Extended Data Figure 1:**
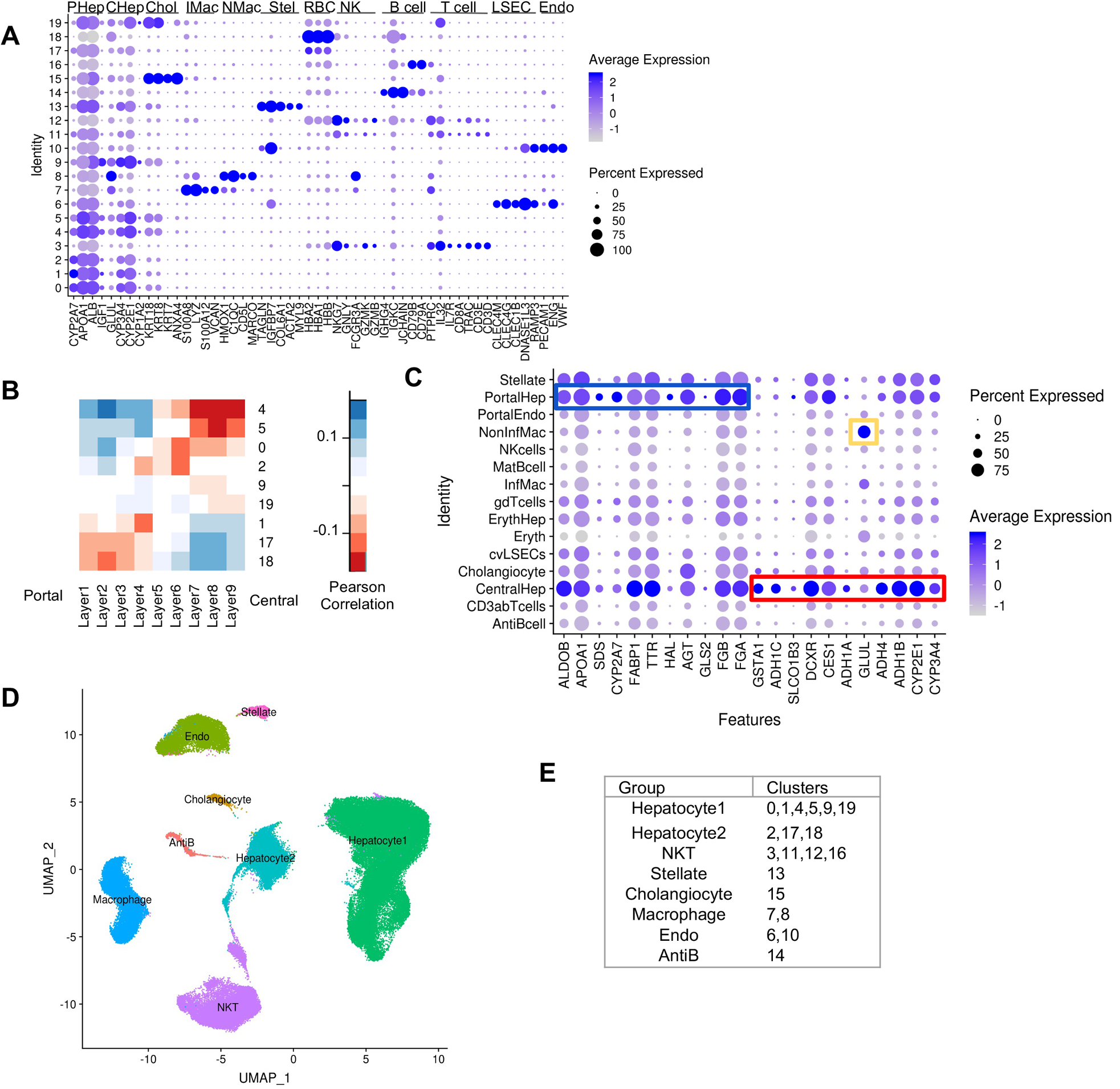
Annotation of Coarse-level map. (A) Marker gene expression used to manually annotate 20 healthy liver clusters. (B) Pearson correlation between human hepatocyte clusters and zonation layers described in mouse (Halpern et al. 2017)). (C) Mouse hepatocyte zonation markers expressed in the annotated human map. (D & E) The map was divided into 8 major subgroups for subclustering.

**Extended Data Figure 2:**
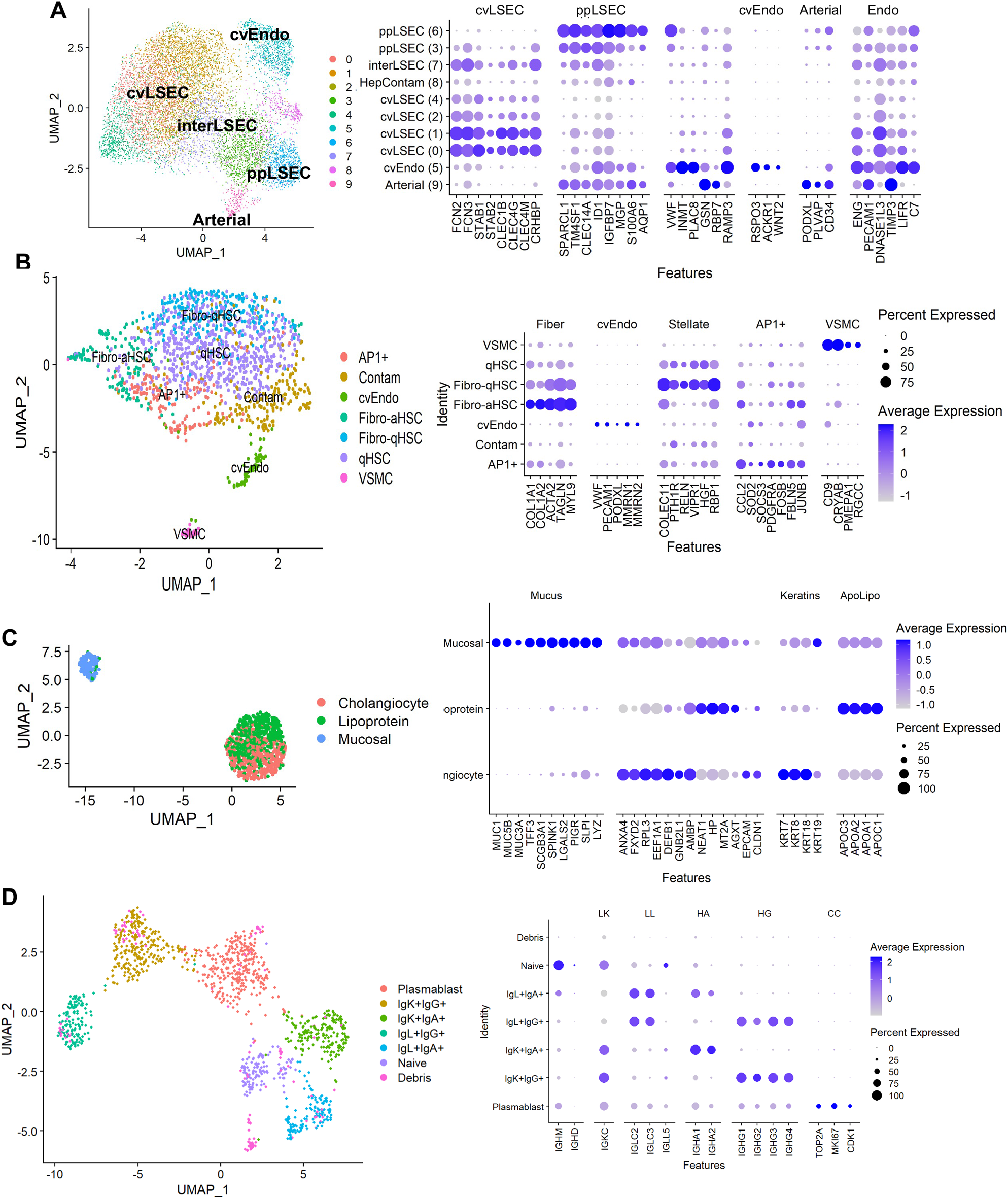
Subclustering of other populations of the Healthy Liver Map. (A) Endothelial cells. (B) Stellate / Mesenchymal cells (C) Cholangiocytes (D) Antibody secreting B cells. Left panels show UMAP of subclusters, Right panel shows top marker genes for each cluster.

**Extended Data Figure 3:**
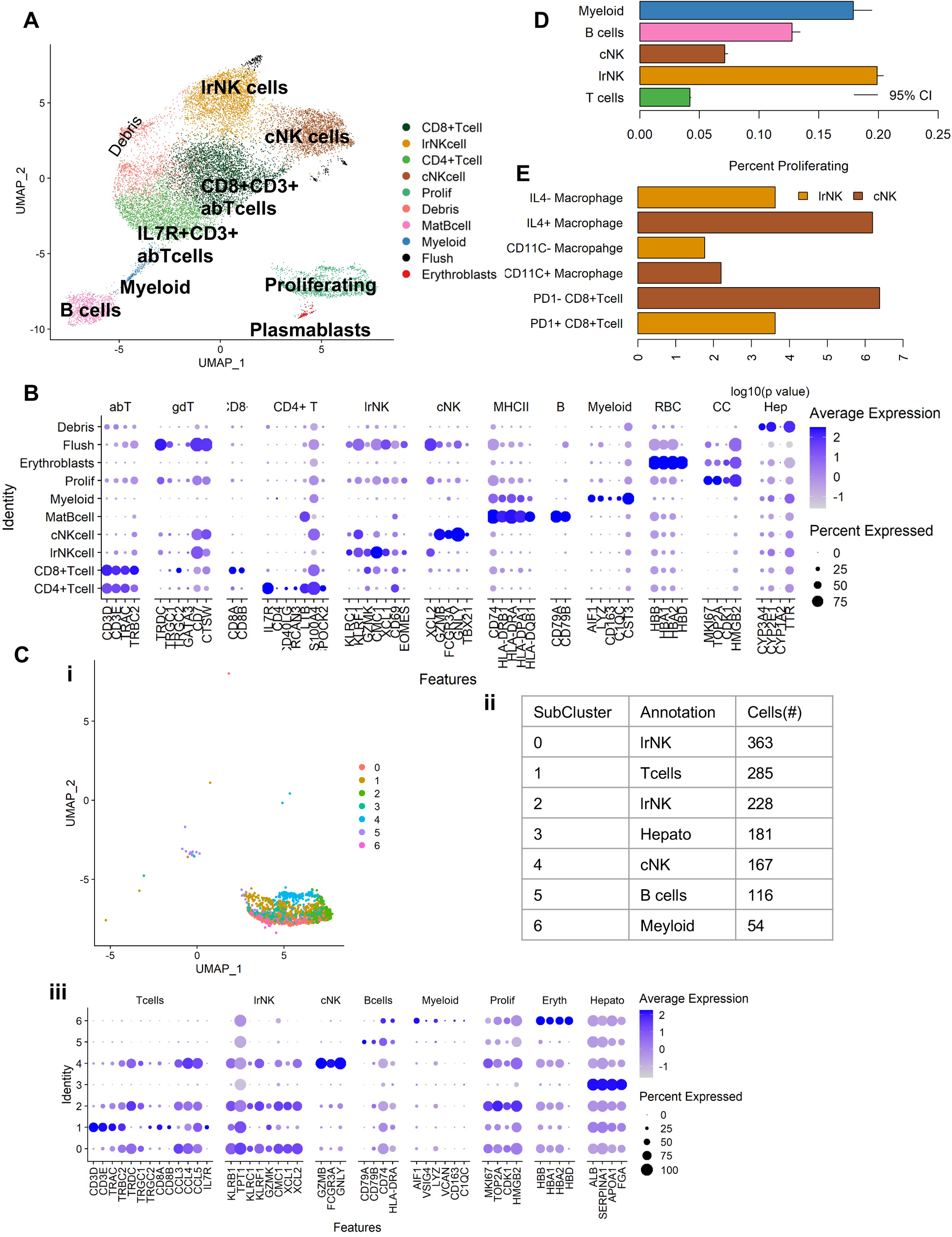
Subclustering of lymphocyte-like cells of the Healthy Map. (A) UMAP of subclustered lymphocyte-like cells. (B) Marker genes of lymphocyte subtypes. (C i-iii) Subclustering of the proliferating cluster and annotation using the same markers as B. (D) Proportion of proliferating cells for each lymphocyte group based on C. (E) Differential pathway enrichments between cNK and lrNK cells from GSEA.

**Extended Data Figure 4:**
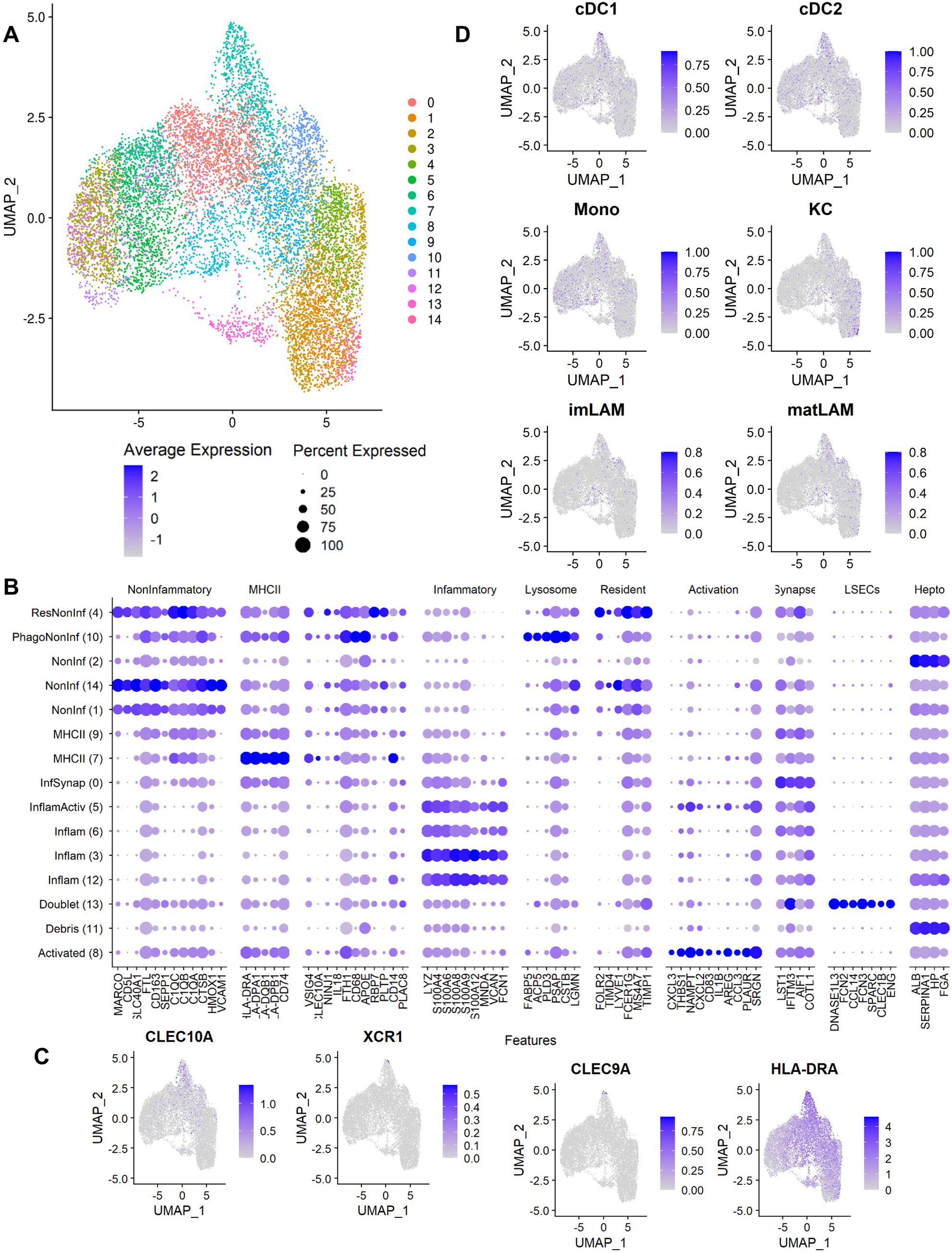
Subclustering of Macrophages. (A) UMAP of integrated subclustered macrophages (B) Top marker genes for each cluster used to manually annotate the macrophage map. (C) Expression of conventional dendritic cell (cDC) markers. (D) Average expression of macrophage gene signatures from Guilliams et al 2022.

**Extended Data Figure 5:**
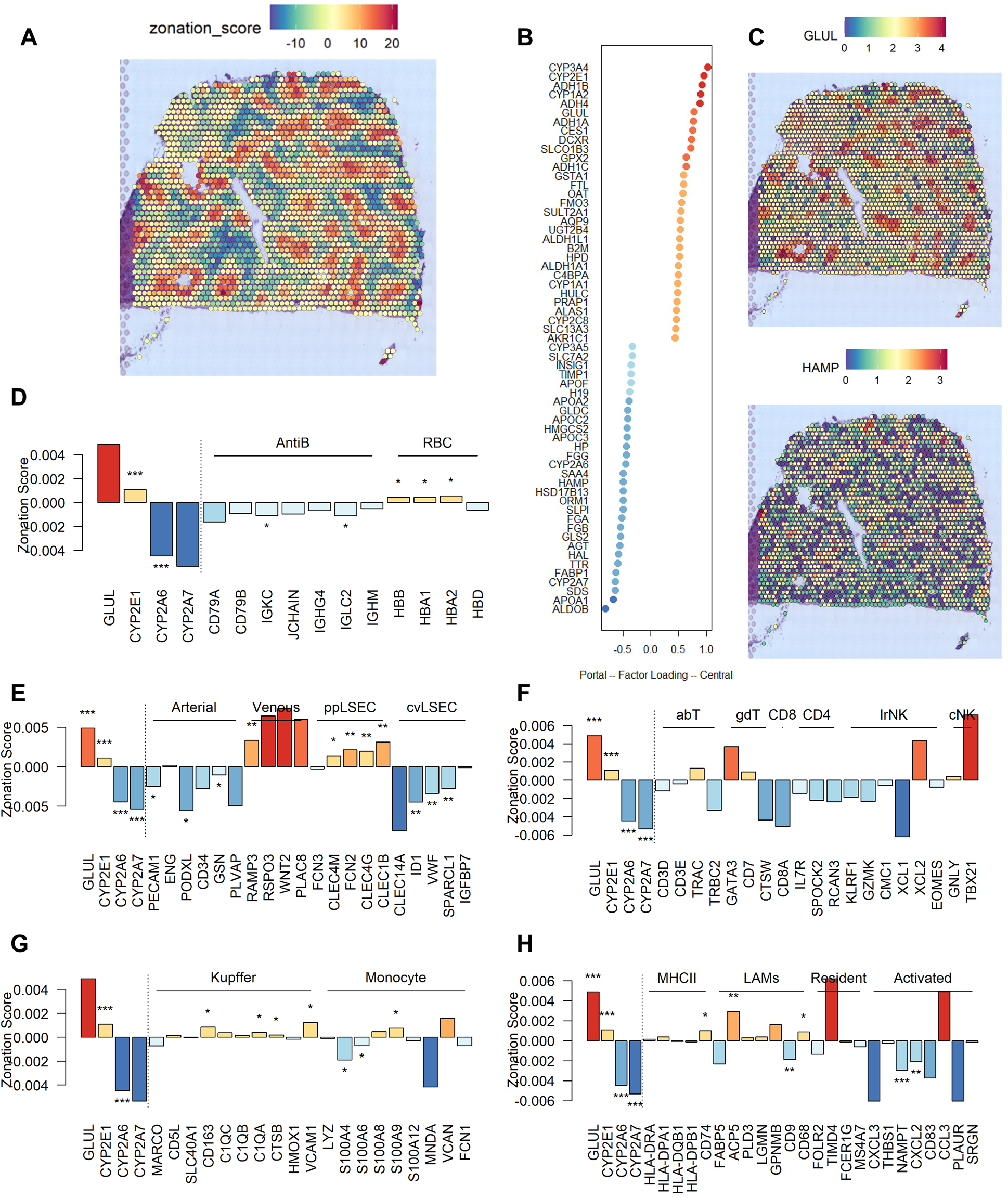
Zonation of cell types in healthy liver. (A) First varimax principal component captures hepatocyte zonation in healthy liver spatial transcriptomics. (B) Top most contributing genes to the zonation associated component. (C) Example gene expression of a pericentral (GLUL) and periportal (HAMP) gene. (D - H) zonation score of individual marker genes for each of the major cell-types identified in the healthy map. Red = pericentral, blue = periportal.

**Extended Data Figure 6:**
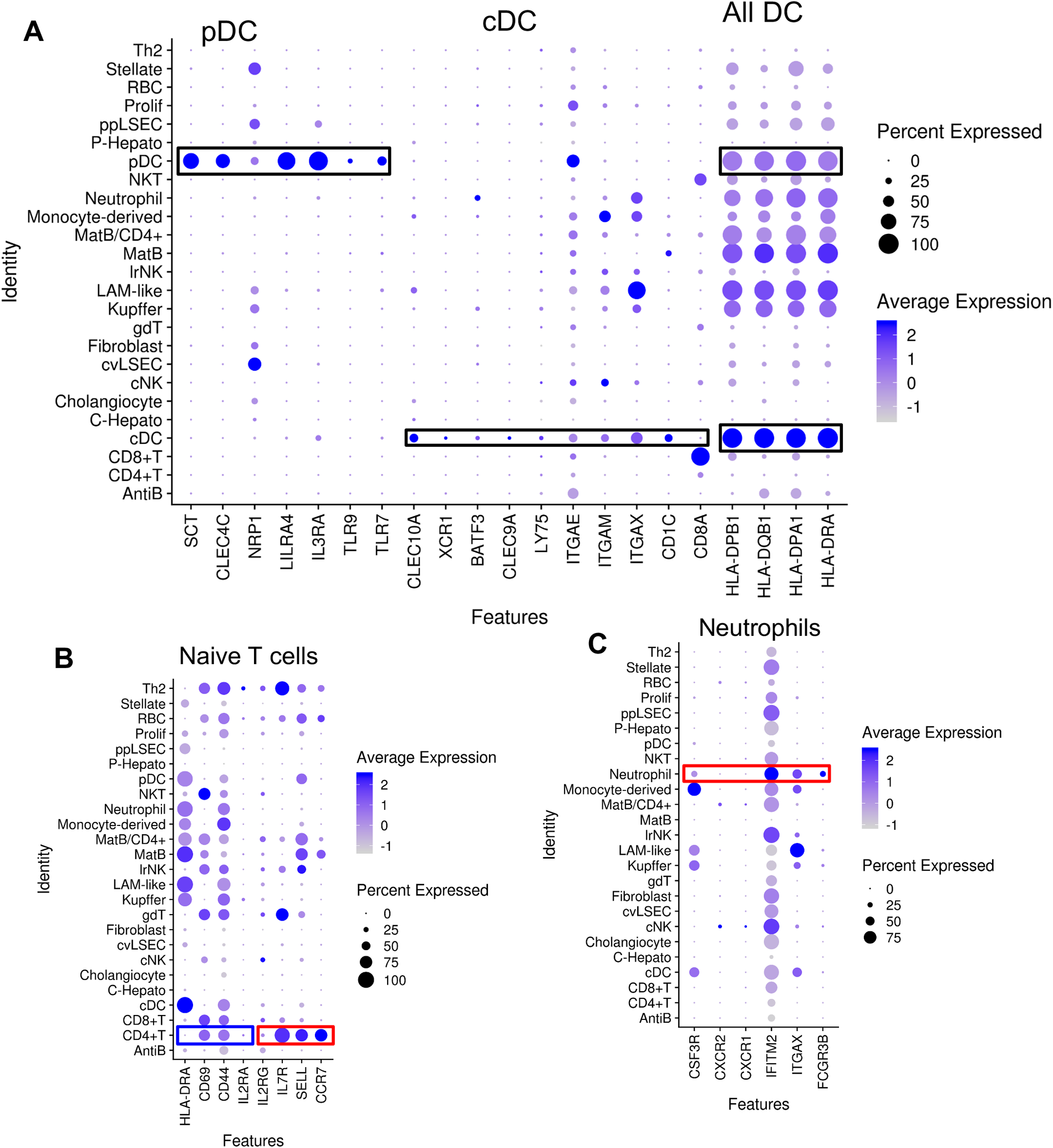
Annotation of PSC Immune cells. (A) Markers of plasmacytoid dendritic cells (pDC), and conventional dendritic cells (cDC). (B) Markers of Naive T cells, red = upregulated in Naive T cells, Blue = downregulated in Naive T cells. (C) Neutrophil markers, note that only FCGR3B is unique to neutrophils.

**Extended Data Figure 7:**
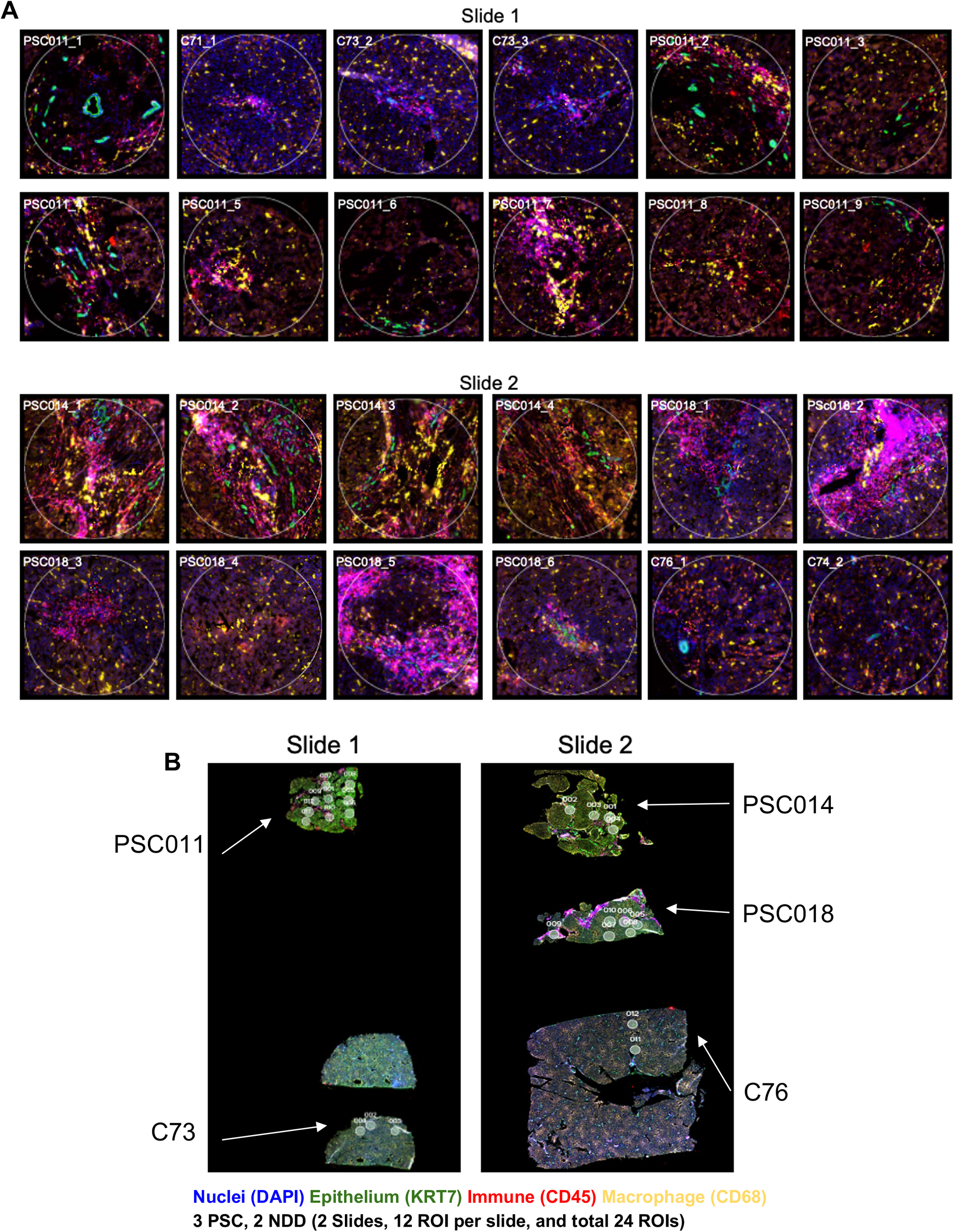
24 Nanostring Regions of Interest (ROIs). Images from the 24 regions of interest (ROIs) selected from the Nanostring GeoMx Digital Spatial Profiling platform. (A) PSC and NDD regions of interest in Slide 1 (PSC011_1-9, C71_1-4) and Slide 2 (PSC014_1-4, PSC018_1-6, C76_1-2). (B) Entire scanned Slide 1 (PSC011, C73) and Slide 2 (PSC014, PSC018, C76). White circles are 660um in diameter.

**Extended Data Figure 8:**
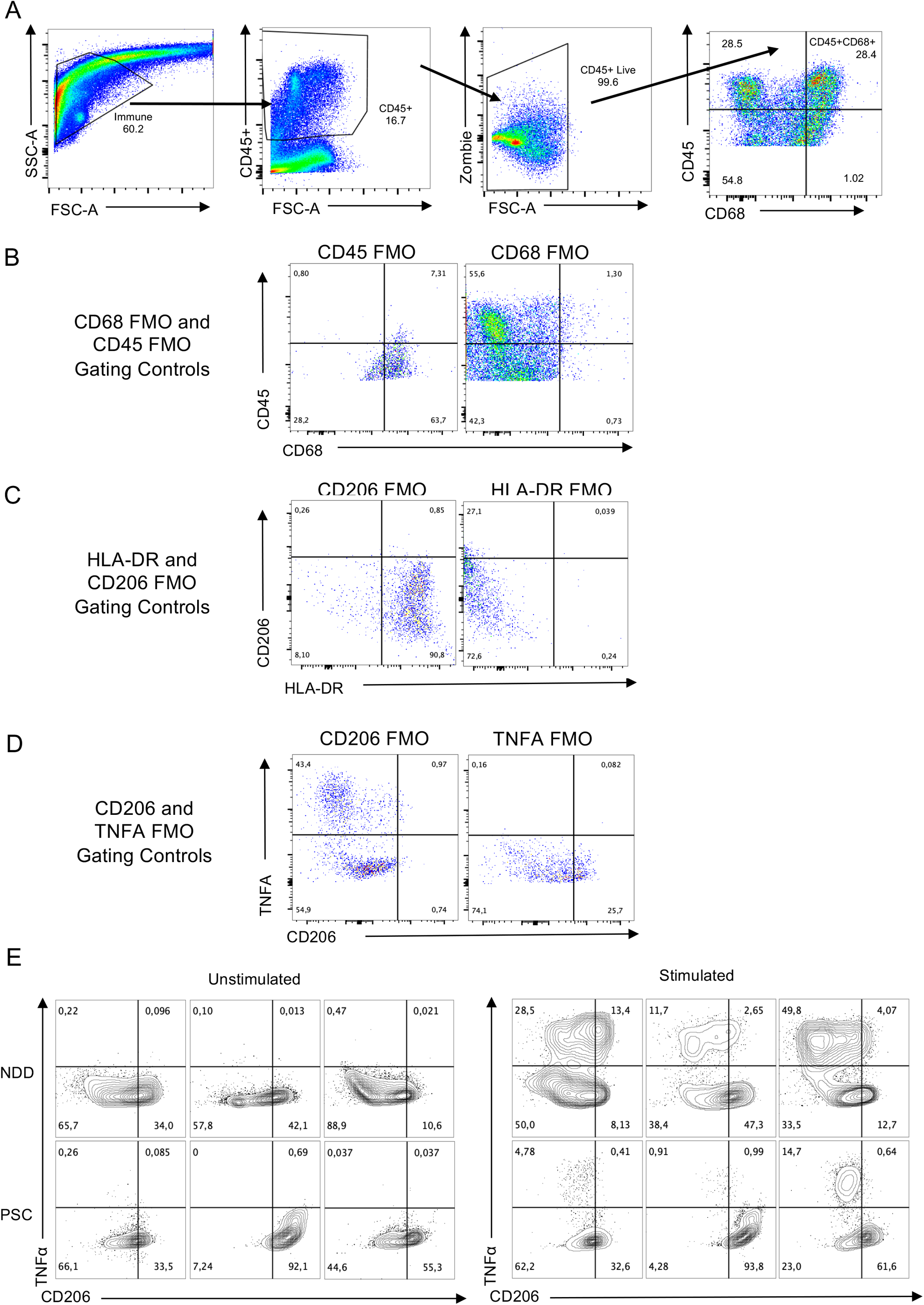
Gating Strategies and FMOs (fluorescence minus one) for flow cytometry immunophenotyping and intracellular cytokine staining of NDD and PSC TLH. A) Representative gating strategy of NDD sample. Gating on the immune fraction in a FSC-A vs SSC-A plot, followed by a gate include CD45+ cells in an FSC-A vs CD45 plot, followed by live cells based on a live/dead Zombie stain in an FSC-A vs Zombie stain, and then gating on CD45+CD68+ cells for macrophage analysis. B) CD45 and CD68 FMO plots used for the gating of CD68+CD45+ cells. C) CD206 and HLA-DR FMOs used to gate on CD206+ and HLA-DR+ cells. C) CD206 and TNFa FMOs used to gate on CD206+TNFa+ cells. E) CD206 and TNFa expression in CD68+CD45+ cells from unstimulated and stimulated NDD and PSC samples (Figure 6).

**Extended Data Figure 9:**
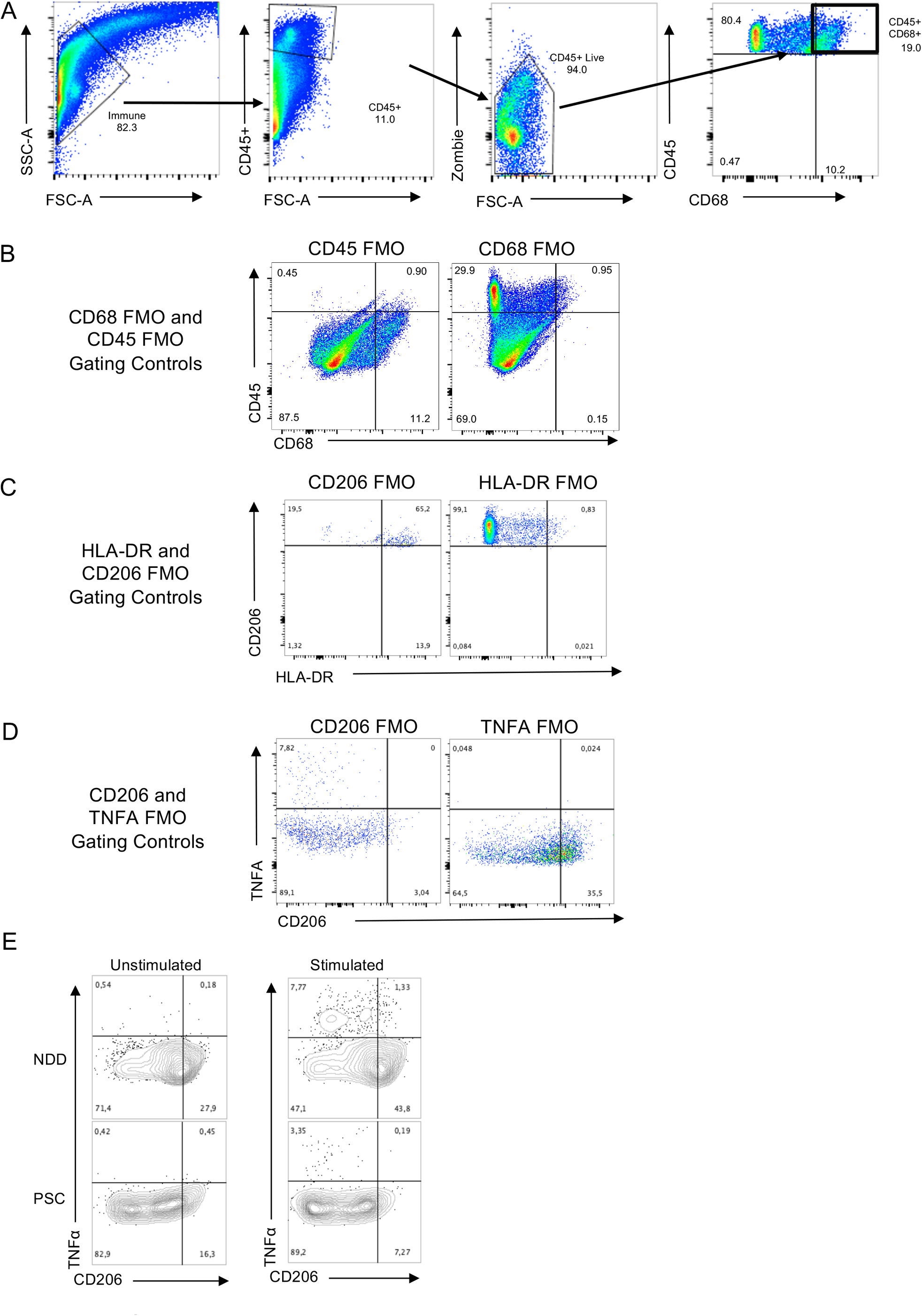
Gating Strategies and FMOs (fluorescence minus one) for flow cytometry immunophenotyping and intracellular cytokine staining of NDD and PSC TLH. A) Representative gating strategy of NDD sample. Gating on the immune fraction in a FSC-A vs SSC-A plot, followed by a gate include CD45+ cells in an FSC-A vs CD45 plot, followed by live cells based on a live/dead Zombie stain in an FSC-A vs Zombie stain, and then gating on CD45+CD68+ cells for macrophage analysis. B) CD45 and CD68 FMO plots used for the gating of CD68+CD45+ cells. C) CD206 and HLA-DR FMOs used to gate on CD206+ and HLA-DR+ cells. C) CD206 and TNFa FMOs used to gate on CD206+TNFa+ cells. E) CD206 and TNFa expression in CD68+CD45+ cells from unstimulated and stimulated NDD and PSC samples (Figure 6).

**Extended Data Figure 10.**
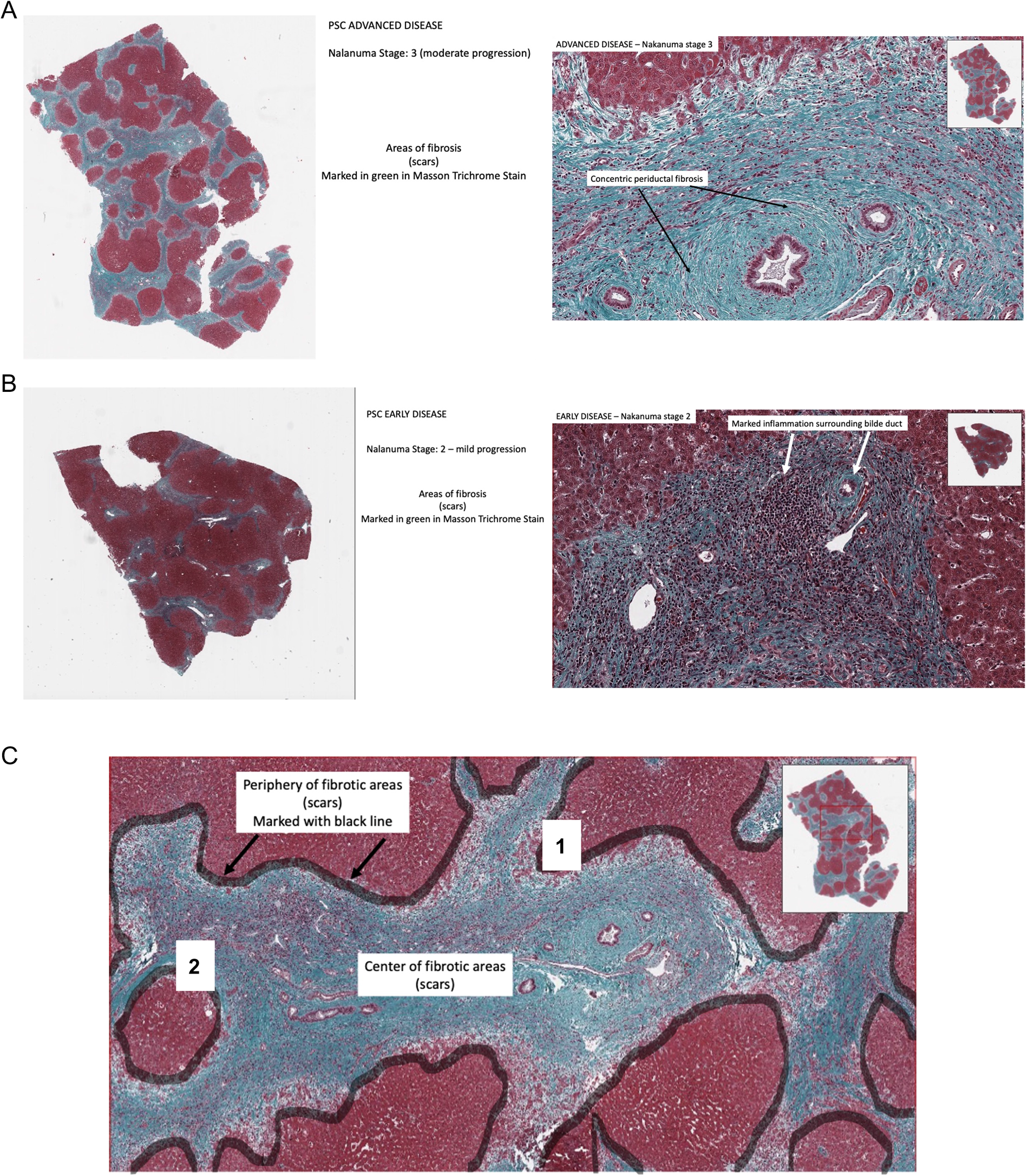
Disease stage according to fibrosis was defined by the Nakanuma score and stage (see reference) assessing the following components: Fibrosis and Bile Duct Loss. A) PSC advanced disease: Masson’s Trichrome stain assessed the extent of fibrosis and identified areas of scarring (stained in green) throughout the liver parenchyma and around individual bile ducts (concentric periductal fibrosis). B) PSC early disease: Early disease PSC livers showed less areas of fibrosis and prominent mononuclear cell inflammation surrounding individual bile ducts. C) Annotation of fibrotic areas (scars) within the liver parenchyma of explanted PSC tissue based on Masson’s Trichrome stain: The fibrotic areas (highlighted in green in Masson Trichrome Stain) are divided into a central and peripheral zone. The peripheral zone is defined as either the interphase between periportal and zone 1 hepatocytes (1) or the periseptal zone of regenerative nodules (2) in cirrhotic liver parenchyma.

**Extended Data Figure 11.**
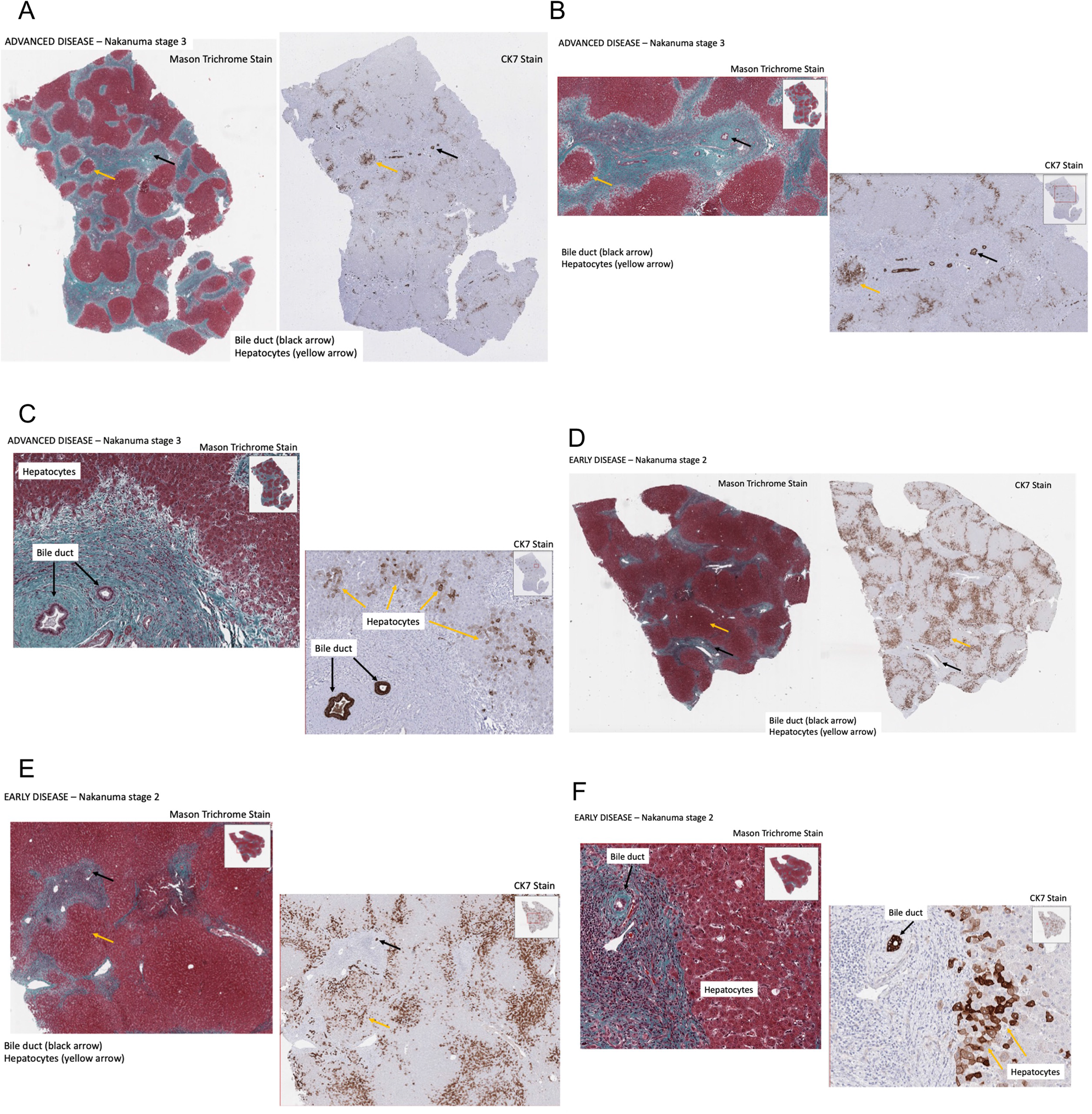
Cytokeratin 7 (CK7) immunohistochemical stain highlights hepatocytes with biliary metaplasia as a feature of chronic biliary disease within fibrotic areas (scars) in PSC explant livers. Bile duct epithelium is marked by a black arrow and metaplastic hepatocytes are marked by a yellow arrow, mainly at the periphery of fibrotic areas (scars). A) PSC advanced disease: Low power view of Masson’s Trichrome Stain and immunohistochemical stain for CK7. B) PSC advanced disease: Intermediate power view of Masson Trichrome Stain and immunohistochemical stain for CK7. C) PSC advanced disease: High power view of Masson Trichrome Stain and immunohistochemical stain for CK7. D) PSC early disease: Low power view of Masson Trichrome Stain and immunohistochemical stain for CK7. E) PSC early disease: Intermediate power view of Masson Trichrome Stain and immunohistochemical stain for CK7. F) PSC early disease: High power view of Masson Trichrome Stain and immunohistochemical stain for CK7.

**Extended Data Figure 12:**
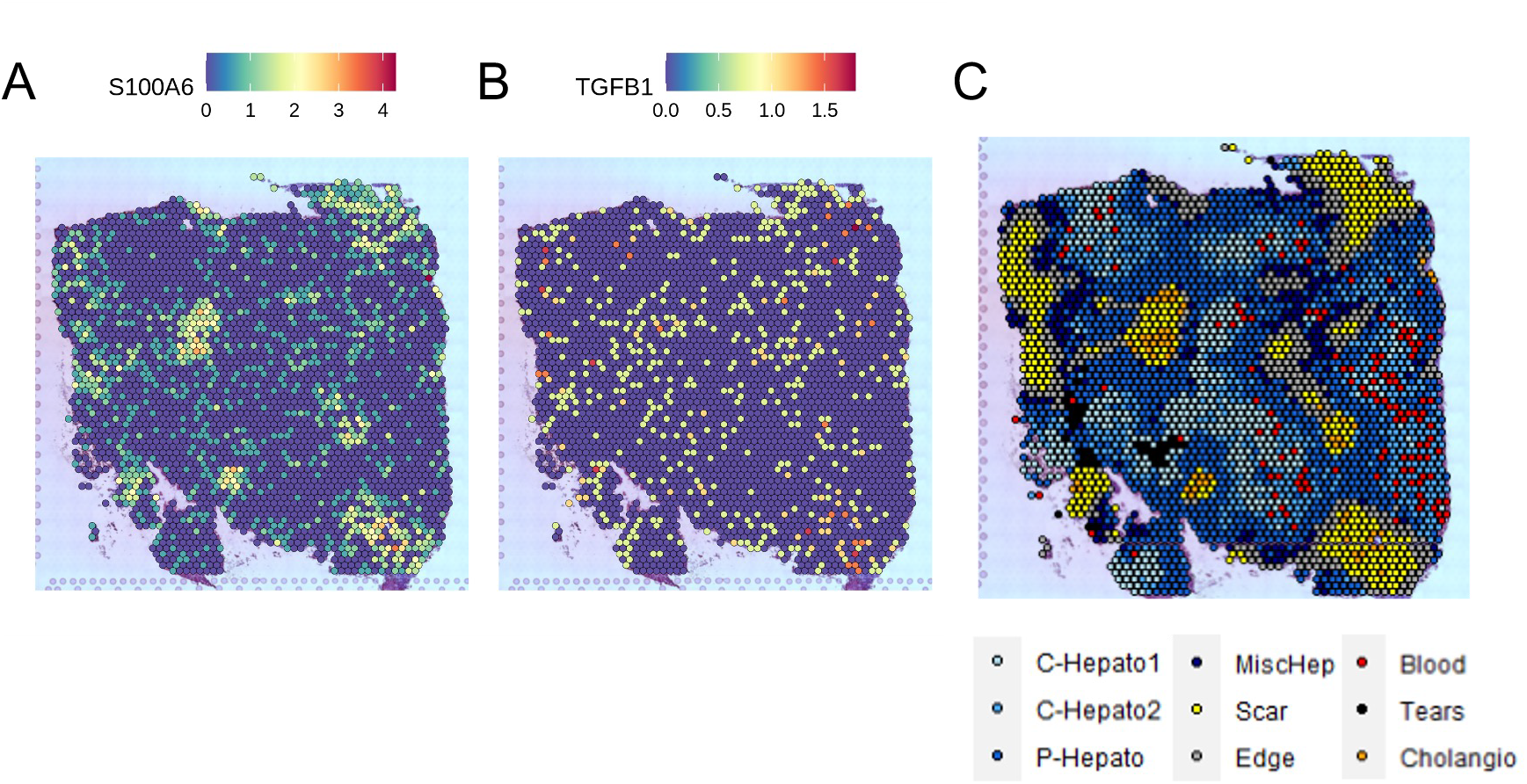
TGFβ expressed in large scars surrounded by transitioning hepatocytes. (A) Expression of the Monocyte-like Macrophage marker S100A6. (B) Expression of TGFβ (C) Annotated clusters in PSC, “Edges” contain transitioning hepatocytes.

## Notes

### Competing Interest Statement

The authors have declared no competing interest.

